# PIN FORMED 2 facilitates the transport of Arsenite in *Arabidopsis thaliana*

**DOI:** 10.1101/710160

**Authors:** Mohammad Arif Ashraf, Kana Umetsu, Olena Ponomarenko, Michiko Saito, Mohammad Aslam, Olga Antipova, Natalia Dolgova, Cheyenne D. Kiani, Susan Nezhati, Keitaro Tanoi, Katsuyuki Minegishi, Kotaro Nagatsu, Takehiro Kamiya, Toru Fujiwara, Christian Luschnig, Karen Tanino, Ingrid Pickering, Graham N. George, Abidur Rahman

**Affiliations:** United Graduate School of Agricultural Sciences, Iwate University, Morioka, Iwate Japan; Department of Plant Bio Sciences, Faculty of Agriculture, Iwate University, Morioka Iwate, Japan; Molecular and Environmental Science Research Group, Department of Geological Sciences, University of Saskatchewan, Saskatoon, Saskatchewan, Canada; Argonne National Lab, Advanced Photon Source, XSD-MIC, Argonne, USA; Isotope Facility for Agricultural Education and Research, Graduate School of Agricultural and Life Sciences, The University of Tokyo, Bunkyo-ku, Tokyo, Japan; Department of Radiopharmaceuticals Development, National Institute of Radiological Sciences, National Institutes for Quantum and Radiological Science and Technology, Inage, Chiba, Japan; Department of Applied Biological Chemistry, Graduate School of Agricultural and Life Sciences, The University of Tokyo, Bunkyo-ku, Tokyo, Japan; Department of Applied Genetics and Cell Biology, University of Natural Resources and Life Sciences, Vienna (BOKU), Muthgasse 18, 1180, Wien, Austria; Department of Plant Sciences, College of Agriculture and Bioresources, University of Saskatchewan, Saskatoon, Canada; Agri-Innovation Center, Iwate University, Morioka, Iwate, Japan

**Keywords:** auxin, arsenite, PIN2, trafficking, transport

## Abstract

Arsenic contamination is a major environmental issue as it may lead to serious health hazard. Reduced trivalent form of inorganic arsenic, arsenite, is in general more toxic to plants compared with the fully oxidized pentavalent arsenate. The uptake of arsenite in plants has been shown to be mediated through a large subfamily of plant aquaglyceroporins, nodulin 26-like intrinsic proteins (NIPs). However, the efflux mechanisms, as well as the mechanism of arsenite-induced root growth inhibition, remain poorly understood. Using molecular physiology, synchrotron imaging, and root transport assay approaches, we show that the cellular transport of trivalent arsenicals in *Arabidopsis thaliana* is strongly modulated by PIN FORMED 2 (PIN2) auxin efflux transporter. Direct transport assay using radioactive arsenite, X-ray fluorescence imaging (XFI) coupled with X-ray absorption spectroscopy (XAS), and ICP-MS analysis revealed that *pin2* plants accumulate higher concentrations of arsenite in root compared to wild-type. At the cellular level, arsenite specifically targets intracellular cycling of PIN2 and thereby alters the cellular auxin homeostasis. Consistently, loss of PIN2 results in aresenite hypersensitivity in root. XFI coupled with XAS further revealed that loss of PIN2 results in specific accumulation of arsenical species, but not the other metals like iron, zinc or calcium in the root tip. Collectively, these results demonstrate that PIN2 serves as a putative transporter of arsenical species *in planta*.

## Introduction

It is estimated that more than 140 million people worldwide are affected by the elevated levels of arsenic (As) in drinking water (WHO 2018). Plants readily accumulate arsenic from contaminated soils and irrigation water, resulting in arsenic exposure to population, especially in arsenicosis-affected areas where irrigated crops are the main staple of the diet (Meharg and Zhao, 2012). Breeding of resistant crop varieties with low accumulation of arsenic requires understanding of complex molecular interactions involved in arsenic uptake, biotransformation, compartmentalisation, and extrusion mechanisms in plants.

Arsenic has a range of oxidation states from −3 to +5 and forms a large variety of organic and inorganic compounds. In natural aquifers, arsenic oxyanions, pentavalent arsenates iAs(V) and reduced trivalent arsenites iAs(III), are predominant inorganic species in aerobic and anaerobic environments, respectively. Trivalent arsenicals are regarded to be more toxic compared to their pentavalent analogues as reviewed in. The toxicity of trivalent arsenicals is connected to their propensity of binding to sulfhydryl groups of proteins resulting in disruption of redox processes and the metabolism of a cell as a whole. In a broad range of pH (pKa=9.2), solvated iAs(III) species is present as uncharged non-dissociated As(OH)_3_ polyol molecule, which structurally and chemically resembles glycerol (Ravenscroft et al., 2009; Yang et al., 2012). Like microorganisms and mammalian cells, higher plants also use aquaglyceroporin proteins to facilitate arsenite entry in plant root cells. It was shown that three of the five plant aquaporin subfamilies, such as nodulin 26-like intrinsic proteins (NIP), plasma membrane (PIP) and tonoplast intrinsic proteins (TIP) are involved in the uptake and translocation of iAs(III) species and methylated organic arsenic metabolites, in plant cells and tonoplasts, respectively. Many of aquaporins demonstrate bidirectional transport properties for arsenic species, so their action can result in efflux of the arsenic to environment (Maciaszczyk-Dziubinska et al., 2012; Xu et al., 2015).

In aerobic conditions, the plant phosphate transporters are the main channel of inorganic As(V) species’ uptake into the cell, where they interfere with processes of oxidative phosphorylation (Shen et al., 2013; Latowski et al., 2018). Inside the plant cell, As(V) is readily reduced to As(III) species with the help of arsenate reductases, of ACR2 and HAC1 (Ellis et al., 2006; Salt, 2017). As-hyperaccumulating ferns from the Pteris genus make use of ACR3-like transporters, absent in angiosperms, to accumulate inorganic arsenite in vacuoles of shoot tissues and gametophytes, possibly as a defence against herbivores (Indriolo et al., 2010). In angiosperms, however, one of the main detoxification mechanisms is formation of As(III) complexes with sulfhydryl (−SH) groups in glutathione and cysteine-rich polypeptides, plant phytochelatines (PCs) and metallothioneins, which are sequestrated in the vacuoles with the help of ABCC transporters (Pickering et al., 2000; Shen et al., 2013; Song et al., 2014). Reduction of As(V) to As(III) was also shown to facilitate excretion of arsenicals back to external medium. It has been found that non-hyperaccumulating species store a majority of As(III)-thiolated species in root vacuoles (Latowski et al., 2018).

Arsenite loading and transport into root vascular system are found to be modulated by several transporters. In Arabidopsis, AtNIP1;1 and AtNIP3;1, and in rice, OsNIP2;1 (Lsi1) have been characterized as major arsenite uptake carriers (Ma et al., 2008; Kamiya et al., 2009; Xu et al., 2015). The efflux of arsenite from the exodermis and endodermis cells to xylem in rice root is suggested to be largely regulated by a silicon efflux carrier Lsi2 (Ma et al., 2008). Recently, involvement of auxin transporter, AUX1 in arsenite response has been shown. The plant tolerance to arsenite is linked to AUX1 mediated auxin transport and reactive oxygen species (ROS)◻mediated signaling (Krishnamurthy and Rathinasabapathi, 2013). However, the role of auxin transporters in arsenite transport was not investigated.

The family of PIN-FORMED (PINs) are the major transporters that facilitates the cellular auxin redistribution and homeostasis that directly affects plant growth and development under both optimal and stressed conditions (Okada et al., 1991; Luschnig et al., 1998; Shibasaki et al., 2009; Hanzawa et al., 2013; Wu et al., 2015; Ashraf and Rahman, 2019). The analysis of structure and function of PINs’ family of auxin efflux carriers places them as a part of bile/arsenite/riboflavin transporter (BART) superfamily of secondary transporters and signalling proteins (http://www.tcdb.org/search/result.php?tc=2.A.69#ref9696, Mansour et al., 2007). Notably, members of BART include Arc3 family of arsenical resistance bacterial proteins transporting As(III) and Sb(III) (Maciaszczyk-Dziubinska et al., 2012).

Many proteins are not confined to only one function. For example, the LSi2, plasma membrane silicic acid efflux pump, is also a member of Arsenite-Antimonite (ArsB) efflux family, and serves as an arsenite efflux transporter in plants (Ma et al., 2008). With that, LSi2 shows 18% identity to the to the *Escherichia Coli* (E.coli) efflux transporter ArsB (Ma et al., 2008). Although these two transporters from two different species show very low identity, at cellular level they execute a similar function raising the possibility that other plant transporters homologous to ArsB may show a similar functionality. In fact, it was previously reported that portions of auxin efflux facilitator (EIR1)/PIN FORMED 2 (PIN2) show 35-40% similarity to *E.coli* efflux carrier ArsB, and to SbmA, an *E.coli* integral membrane protein, which is required for the uptake of the antibiotic Microcin 25 (Luschnig et al., 1998).

Although PIN2 shows a higher homology to ArsB compared to Lsi2, no effort has been made to characterize whether PIN2 plays any functional role in arsenite transport. Using physiology, molecular and cell biology, high resolution synchrotron imaging, and direct transport assay approaches, we tried to decipher the role of PINs in regulating arsenic response. Our results demonstrate that 1) arsenite, but not the arsenate response in Arabidopsis root is regulated by PIN2; 2) arsenite alters the intracellular auxin homeostasis through selective modulation of PIN2 trafficking, 3) Loss of PIN2 specifically affects the accumulation of arsenical species, but not the other metals like zinc, iron or calcium in the Arabidopsis root tip, and 4) PIN2 facilitates the transport of trivalent arsenical species and functions as a putative efflux transporter for arsenite metabolites *in planta*.

## Results

### Homology of plant auxin efflux carriers and selected arsenite transporters

Previously it was reported that portions of PIN2 show 35% - 40% identity to the bacterial transporter ArsB (Luschnig et al., 1998). We reassessed the homology of PIN proteins with different arsenite transporters using a bioinformatics approach. Plasma membrane localized PIN proteins, which function as intracellular auxin efflux carriers, all have a similar structure, with two hydrophobic domains, consisting of about 5 transmembrane helixes each, separated by a central intracellular hydrophilic domain. Among the 8 annotated PIN proteins in Arabidopsis, PIN1, PIN2, PIN3, PIN4, and PIN7 reside in the plasma membrane, while PIN5 and PIN8 are localized in the endoplasmic reticulum (ER). Recent research showed a complex behavior and localization of PIN6 both in ER and plasma membrane (Simon et al., 2016).

The cladogram of the plasma membrane localized PIN proteins places PIN1 and PIN2 in one clade, and PIN3, PIN4 and PIN7 in another clade, where PIN3 and PIN7 are closely associated because of their high homology (Supplemental Figure 1). In general, the homology among plasma membrane residing PIN proteins ranges from 60% to 90% (Supplemental Figure 2 and Supplemental Table 1). Multiple sequence alignments of PIN1, PIN2 and PIN3 proteins against various arsenite transporters revealed that they show approximately 25% homology with bacterial transporter ArsB, but lower homology with Lsi2 (Table 1, Supplemental Figure 3 and Supplemental Table 2). AtPINs also showed 18% homology to arsenite transporters Acr3 from yeast *S. cerevisiae* (Ghosh et al., 2002), and arsenic hyperaccumulator fern *Pteris vittata*^11^. Compared with Lsi2, AtPINs show higher homology to all the known arsenite transporters (Table 1, Supplemental Table 3).

**Table 1:** Homology of plant auxin efflux carrier (PINs) and arsenite transporters

|  | SsAcr3 | PvAcr3 | arsB | OsLsi2 | AtPIN2 | AtPIN1 | AtPIN3 |
| --- | --- | --- | --- | --- | --- | --- | --- |
| SsAcr3 | 100.00 | 40.16 | 8.22 | 6.44 | 16.67 | 18.01 | 17.28 |
| PvAcr3 | 40.16 | 100.00 | 13.99 | 6.86 | 17.65 | 18.45 | 18.13 |
| arsB | 8.22 | 13.99 | 100.00 | 12.15 | 24.51 | 24.44 | 24.86 |
| OsLsi2 | 6.44 | 6.86 | 12.15 | 100 | 10.26 | 9.63 | 8.83 |
| AtPIN2 | 16.67 | 17.65 | 24.51 | 10.26 | 100.00 | 62.88 | 60.88 |
| AtPIN1 | 18.01 | 18.45 | 24.44 | 9.63 | 62.88 | 100.00 | 65.49 |
| AtPIN3 | 17.28 | 18.13 | 24.86 | 8.83 | 60.88 | 65.49 | 100.00 |

Identity matrix of *Saccharomyces cerevisiae* Acr3 (SsAcr3), *Pteris vittata* Acr3 (PvAcr3) and *Escherichia coli* arsenite transporter (arsB), *Oryza sativa* silicon transporter (OsLsi2), and *Arabidopsis thaliana* PINs (AtPIN1, AtPIN2, AtPIN3).

### Loss of PIN2 results in altered response to arsenite

To understand the functional significance of AtPINs homology to bacterial arsenite transporter ArsB, we next investigated the response of selected PIN1, PIN2, PIN3 and PIN4 mutants to both arsenite and arsenate. Root elongation in *A. thaliana* shows a strong response to exogenous arsenite and arsenate, albeit at different concentrations. Time course and dose response assays of root growth in wildtype revealed that approximately 50% inhibition of root elongation can be achieved with 10μM arsenite over 3 days incubation (Figure 1A and 1B). Consistent with previous results (Lee et al., 2003), a much higher concentration of arsenate (1.5mM) was required to achieve similar degree of root elongation inhibition (Supplemental Figure 4).

**Figure 1:**
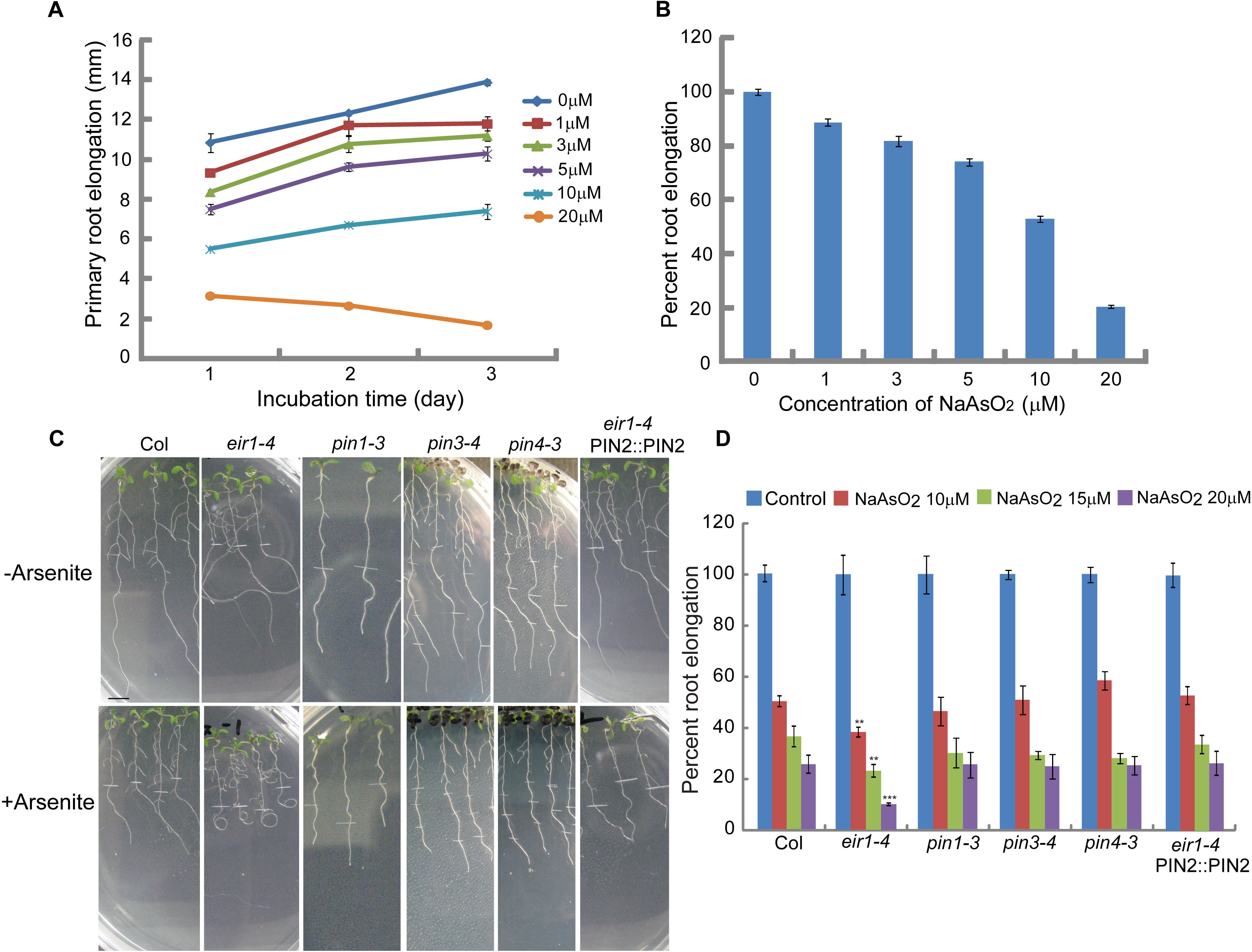
Effect of arsenite on wild-type and mutants root elongation response. Five-day-old light grown wild-type or mutants seedlings were transferred to new agar plates supplemented with or without arsenite and incubated for various time lengths under continuous light. **(A)** Time course of arsenite induced inhibition of root elongation in wild-type. **(B)** Dose response of arsenite for root elongation in wild-type after 3-day incubation. Approximately fifty percent inhibition of root growth was observed at 10 μM arsenite. **(C)** Representative images of root phenotype of wild-type, *pin* mutants and *pin2* complemented line after 10 μM arsenite treatment for 3 days. Bar represents 0.5cm. **(D)** *pin2/eir1-4* mutant shows hypersensitive response to arsenite induced root growth inhibition. Five-day-old Arabidopsis seedlings were subjected to arsenite treatment for 3 days. Compared with wild-type, *eir1-4* showed hypersensitive response to arsenite induced inhibition of root elongation at all concentrations we tested (P < 0.0001), while complemented line of *pin2*, *eir1-4*-PIN2:PIN2 show wild-type like response to arsenite induced root growth inhibition. For data shown in A, B and D vertical bars represent mean ± S.E. of the experimental means from at least five independent experiments (*n* = 5 or more), where experimental means were obtained from 8-10 seedlings per experiment.

In a previous report, it was claimed that both *pin2* and *pin1* mutants were hypersensitive to arsenite-induced root growth inhibition (Krishnamurthy and Rathinasabapathi, 2013). However, in our screening, *pin1*, *pin3* and *pin4* mutants showed a wild-type response to arsenite exposure (Figure 1C and 1D).

Among the membrane residing PIN mutants that we tested for root growth assay, the response of *pin2* to arsenite exposure was the most striking. At all tested concentrations of arsenite, two independent allele of *pin2* mutant, *eir1-4 and eir 1-1* roots showed hypersensitive response to arsenite-induced inhibition of root elongation. Additionally, roots of both alleles exhibited hook-like curling in presence of arsenite (Figs. 1C and D, Supplemental Figure 4). Since both the alleles show essentially similar response, we used more widely used allele *eri1-1* for subsequent experiments. Complementation of *pin2* mutation with genomic PIN2 reverted back both the curling root phenotype and hypersensitive root growth response, confirming that the observed altered response of *pin2*mutant towards arsenite is linked to PIN2 (Figs. 1C and D). Interestingly, overexpression of PIN2 results in increased resistance to arsenite-induced root growth inhibition (Supplemental Figure 4). In contrast, all these mutants showed wild-type like response to arsenate-induced root growth inhibition (Supplemental Figure 5). Collectively, these results strongly suggest that PIN2 is a potential regulator of arsenite response in roots.

### Arsenite alters auxin response in Arabidopsis root through modulating auxin transport

PIN2 is functional for auxin efflux in the lateral root cap (LRC), epidermal and cortex cells, and through its intracellular polarity it maintains a maximal auxin gradient at the root tip, which is an absolute requirement for root gravity response(Rahman et al., 2010). Consistently, loss of PIN2 results in complete agravitropic response in roots(Luschnig et al., 1998). Since *pin2/eir1-1* mutant showed altered response to arsenite, we hypothesized that arsenite may affect the root auxin response. To clarify this possibility, we investigated the effect of arsenite on the root gravity response in wild type roots. Arsenite considerably slows down the gravity response of the wild type roots. However, effect of arsenite on wild type root elongation was statistically insignificant during the gravity response assay period (Figure 2A and 2B), confirming that the arsenite-induced inhibition of gravity response is unlinked to the inhibition of root elongation.

**Figure 2:**
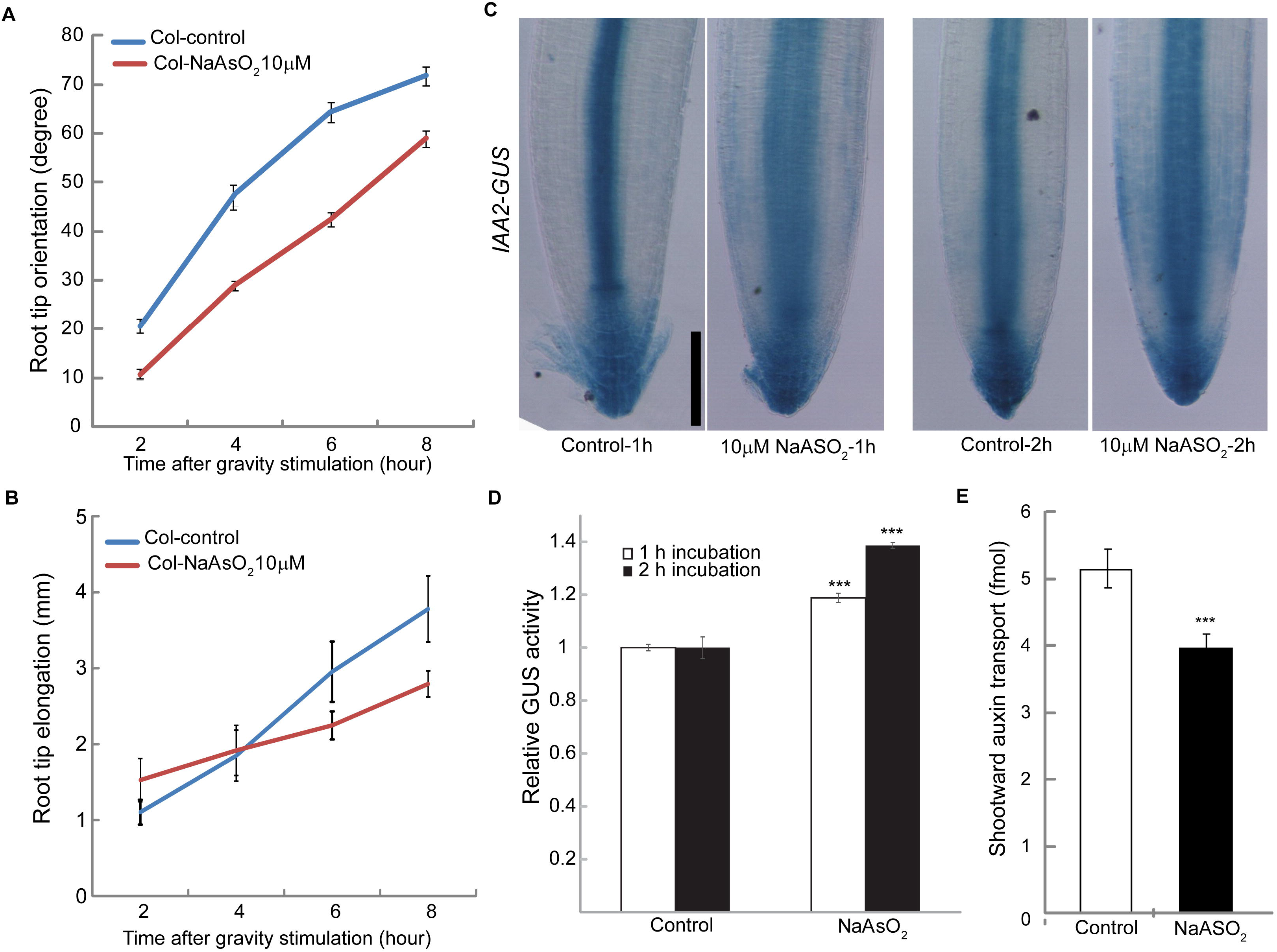
Arsenite inhibits root gravity, alters intracellular auxin response and inhibits auxin transport. **(A)** Effect of Arsenite on root gravity response. For assaying gravitropism, five –day-old light grown seedlings were transferred to arsenite, gravistimuated. Data for root tip orientation was collected for 2,4,6 and 8 h. Vertical bars represent mean ± S.E. of the experimental means from at least five independent experiments (*n* = 5 or more), where experimental means were obtained from 8-10 seedlings per experiment. Arsenite-induced inhibition of root gravity response was significant at all-time points as judged by Student’s *t*-test (P < 0.0001). **(B)** Effect of Arsenite on root elongation during the gravity assay. Arsenite-induced root growth inhibition was insignificant at all-time points as judged by Student’s *t*-test. **(C)** Arsenite alters the intracellular auxin response. Five-day-old light grown *IAA2-GUS* seedlings were treated with 10 μM Arsenite for 1h and 2h respectively. After the Arsenite treatment, GUS staining was performed by incubating the seedlings in GUS staining buffer for 1h at 37°C. Demonstrated images are representative of 15-20 roots obtained from at least three independent experiments. Bar represents 100 μm. **(D)** Quantification of GUS activity obtained from experiment **C**. Vertical bars represent mean ± S.E. Compared with the control treatment, Arsenite-induced increase in GUS activity was highly significant (P < 0.0001) in both time points as judged by Student’s *t*-test. **(E)** Effect of arsenite on shootward auxin transport. Five-day-old light grown seedlings were transferred to new agar plates and subjected to arsenite treatment before transport of ^3^H IAA over 2 h was measured as described in the methods. The experiments were conducted using at least three biological replicates. For each biological replicate, three technical replicates were assayed. (Col-control, *n*= 57; Col-arsenite, *n*=52). Asterisks represent the statistical significance between treatment (*** P < 0.0001). Vertical bars represent mean ± S.E. of the experimental means.

The root gravity response is regulated by the asymmetric distribution of auxin, which is largely dependent on the auxin effluxed by PIN2 (Luschnig et al., 1998; Rahman et al., 2010). To understand whether the cellular auxin homeostasis in the root meristem is altered by arsenite, we monitored the intracellular auxin response using two auxin responsive markers *IAA2-GUS* and DII-VENUS both of which are capable of detecting intracellular auxin distribution at high spatio-temporal resolution (Luschnig et al., 1998; Shibasaki et al., 2009; Band et al., 2012; Brunoud et al., 2012; Hanzawa et al., 2013). Only a brief incubation in arsenite altered the auxin response pattern in root meristem. More GUS staining was observed in arsenite treated roots compared with wild-type, and the response was proportional to the incubation time (Figure 2C and 2D). Similar results were observed with DII-Venus marker line for long term arsenite treatment (Supplemental Figure 6). Interestingly, after 2 h incubation in arsenite, GUS signal started to accumulate in the peripheral cells like epidermis and cortex which typically results from inhibition of auxin transport (Figure 2C, Shibasaki et al., 2009), indicating that arsenite may inhibit auxin transport. Shootward auxin transport (which is largely regulated by PIN2) assay with 3^H^ IAA revealed that arsenite indeed inhibits the shootward auxin transport (Figure 2E).

These results suggest that in addition to other systemic effects likely exhibited by trivalent arsenicals in living cells (Shen et al., 2013), arsenite-modulated PIN2 activity resulted in the altered cellular auxin response and reduced auxin transport.

### Arsenite alters intracellular trafficking of PIN2

To provide a mechanistic explanation of arsenite effect on PIN2, we next investigated expression of PIN2 both at transcriptional and translational levels. The transcript analyses of PIN2 by quantitative real time PCR revealed no significant difference in transcript level under arsenite treatment, suggesting that PIN2 is not under direct transcriptional regulation of arsenite (Supplemental Figure 7). Earlier it has been demonstrated that for proper functioning of PIN2 as an IAA efflux protein, both the polar deployment and intracellular trafficking of PIN2 are required (Luschnig et al., 1998; Laxmi et al., 2008; Shibasaki et al., 2009; Wan et al., 2012; Hanzawa et al., 2013). Moreover, this trafficking process has also been shown to be sensitive to various kinds of stresses (Luschnig et al., 1998; Laxmi et al., 2008; Shibasaki et al., 2009; Wan et al., 2012; Hanzawa et al., 2013). Cellular localization of PIN2, using PIN2-green fluorescent protein (GFP) transgenic seedlings (Xu and Scheres, 2005) revealed that arsenite did not alter the asymmetric localization of PIN2 (Figure 3A, upper panel) but did suppress the trafficking (Figure 3B). In control wild type plants, protein trafficking inhibitor brefeldin A (BFA) resulted in formation of large number of PIN2-positive small bodies in cytosol, supporting the notion of continuous cycling of PIN2 between plasma membrane and endosomal compartments (Figure 3A and Supplemental Figure 8). Importantly, in both short (2h) and long term (3d) arsenite treatments, formation of these small PIN2-positive bodies was drastically reduced (Figure 3A, 3B and Supplemental Figure 8A, 8B).

**Figure 3.**
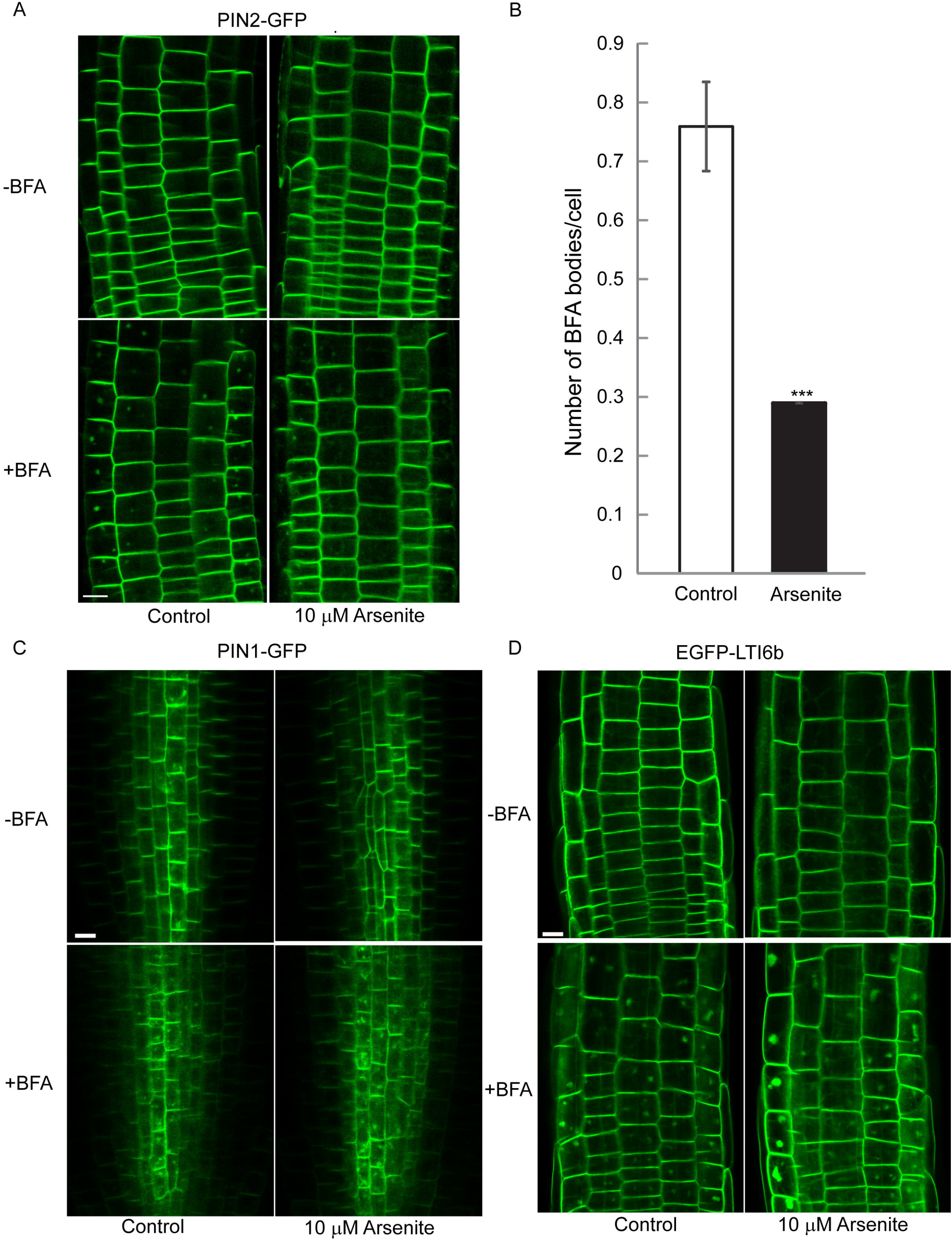
Arsenite specifically affects the intracellular dynamic cycling of PIN2. Five-day-old PIN2::PIN2–GFP, PIN1::PIN1-GFP and EGFP-LTI6b transgenic seedlings were treated with arsenite for 2h. After the incubation, seedlings were treated with 20 μM BFA for 40 min. The images were captured using same confocal setting and are representative of 15-20 roots obtained from at least 4 independent experiments. **(A)** Effect of Arsenite on PIN2 trafficking. Bar represents 10 μm. **(B)** Quantitative analysis of formation of PIN2-BFA body in the transition zone of PIN2::PIN2–GFP transgenic plants in presence or absence of Arsenite. Total number of BFA body and number of cells were counted in the imaged area. Bar graph represents the average number of BFA body formed per cell. Vertical bars represent mean ± S.E. of the experimental means (*n* = 4 or more). Asterisks represent the statistical significance between treatment (*** P < 0.0001). **(C)** Effect of Arsenite on PIN1 and LTI6b trafficking. Note that BFA bodies are formed in presence of Arsenite. Bar represents 10 μm.

To elucidate the specificity of arsenite-induced inhibition of PIN2 trafficking, we investigated its effect on the trafficking of PIN1, a close homologue of PIN2 and LTI6b, a cold-inducible membrane protein, which is trafficked from the plasma membrane to endosomes through a BFA regulated pathway (Kurup et al., 2005; Shibasaki et al., 2009). In both short and long term arsenite treatments, BFA-induced PIN1 and LTI6b bodies were formed (Figure 3C, 3D and Supplemental Figure 8C, 8D), suggesting that arsenite specifically targets PIN2 trafficking.

### Arsenite transport is altered in *pin2* root

In bacteria and yeast, arsenite transport mechanism is extensively studied. In yeast, arsenite is effluxed out of the cells by Acr3, a plasma membrane-localized efflux carrier and some aquaglyceroporines, functioning as bidirectional arsenite transporters (Maciaszczyk-Dziubinska et al., 2012; Yang et al., 2012). In bacteria, arsenite uptake and efflux is passively regulated by the bidirectional aquaglyceroprotein channels and also pumped outside of the cells by ArsB or ArsAB functioning as As(OH)_3_-H^+^ antiporter or ATP-driven extrusion pump, respectively. Some bacteria possess both ArsAB and Acr3 efflux systems (Meharg and Zhao, 2012; Yang et al., 2012).

However, in plants, several aquaglyceroprotein NIPs have been shown to regulate passive, gradient driven arsenite uptake, while only a single protein, rice Lsi2, has been implicated in active regulation of arsenite efflux. Rice Lsi2 is a silicon transporter and shows 18% homology with bacterial arsenite transporter, ArsB (Ma et al., 2008; Meharg and Zhao, 2012). Since PIN2 shows a higher homology with ArsB compared with Lsi2, *pin2/eir1-1* mutant shows hypersensitive response to arsenite-induced root growth inhibition, and arsenite specifically targets the PIN2 trafficking, we hypothesized that PIN2 may mediate arsenite transport in root. To clarify this possibility, we combined ^74,73^As (III) direct transport assay, ICP-MS analysis of arsenic accumulation, and speciation and localization of arsenic in roots using high resolution synchrotron X-ray fluorescence imaging (XFI) analysis coupled with X-ray absorption spectroscopy (XAS).

One of the most reliable methods to show transport activity of a plant protein is direct transport assay *in planta*. For arsenite, this is a challenging issue as it is not commercially available. We solved the problem by developing radioactive arsenite (^74,73^As) by chemical reduction of radioactive arsenic (see supplemental methods for detail explanation). A short term ^74,73^As transport assay (2h) was performed to compare arsenite transport activity in wild type and *pin2*/*eir1-1* mutant plants using radioimaging. Five-day old wild type and *pin2*/*eir1-1* seedlings were incubated in 0.1 and 10 μM ^74,73^As for 2h. The quantification of radioimaged plates revealed a noticeable increase in ^74,73^As activity in *pin2*/*eir1-1* roots compared with wild-type roots (Figure 4A and 4B), suggesting that arsenite transport is impaired in *pin2*/*eir1-1* mutant.

**Figure 4.**
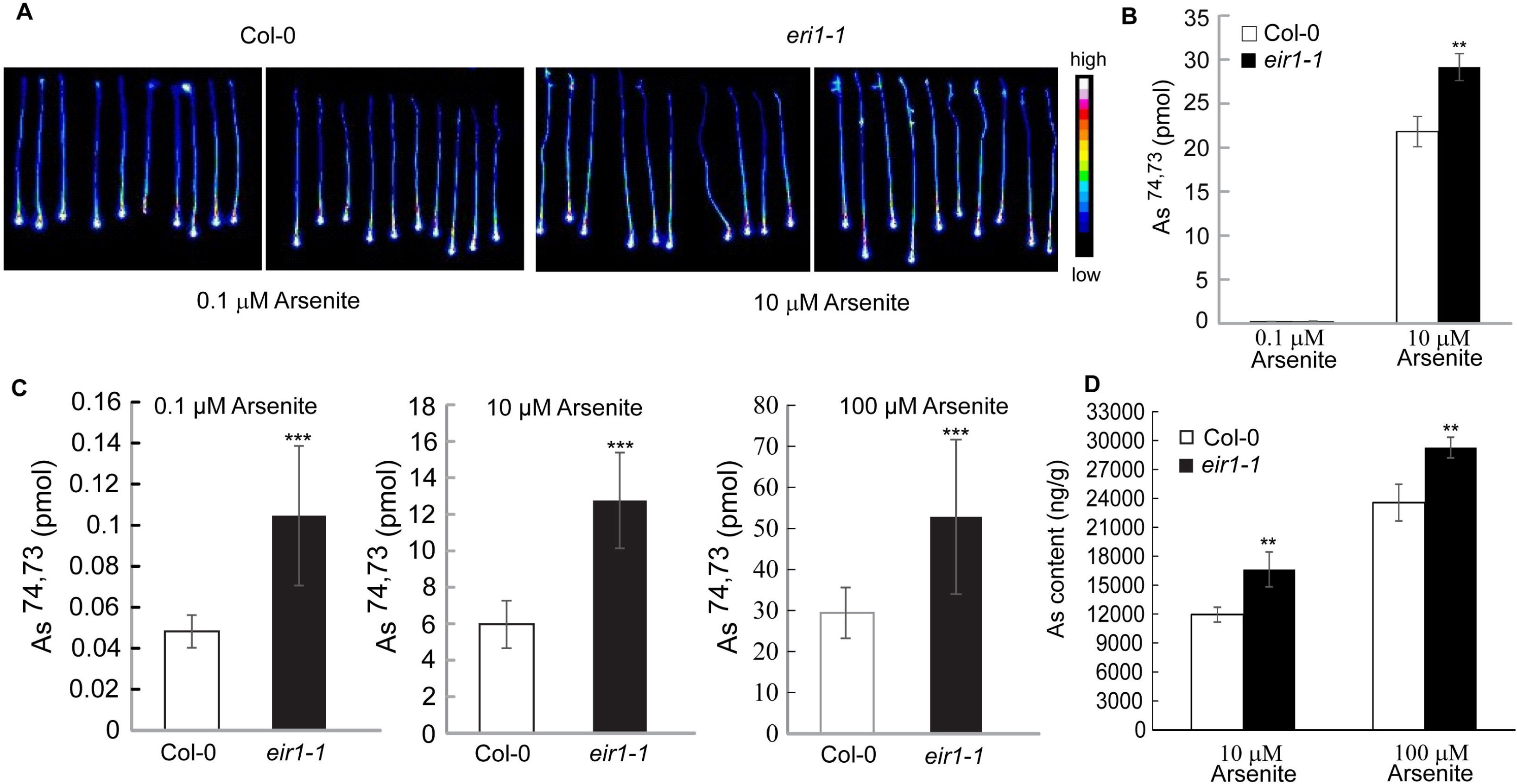
*pin2/eir1-1* shows altered transport and accumulation of Arsenite. Allocation of ^74,73^As in Col-0 and *eir1-1* **(A)**. Five-day-old Col-0 and *eir1-1* roots were incubated in 0.1 μM and 10 μM ^74,73^As for 2h. ^74,73^As radiation was captured by an imaging plate (IP). Images are representative of three independent experiments. **(B)** Quantification of As allocation in root from experiment **A**. The data were obtained from three independent experiments with 10 seedlings in each treatment. Vertical bars mean ±SE. Asterisks represent the statistical significance between treatments as judged by Student’s *t*-test: **P < 0.001. **(C)**Scintillation counting ^74,73^As activity in Col-0 and *eir1-1*. Five-day-old Col-0 and *eir1-1* seedlings were incubated for 2h at 0.1, 10 and 100 μM ^74,73^ As. Whole root was collected after the incubation. ^74,73^ As activity was measured by liquid scintillation counting. The data were obtained from 10 individual roots for each treatment. Vertical bars mean ±SD. Asterisks represent the statistical significance between treatments as judged by Student’s *t*-test: ***P < 0.0001. **(D)** Arsenic content in Col-0 and *eir1-1*. Five-day-old light grown Col-0 and *eir1-1* seedlings were transferred to 10 and 100 μM arsenite solution and incubated for 2h. 5 mm root tip of 20 seedlings for each treatment was used to measure As by ICP-MS. The data were obtained from three independent experiments. Vertical bars mean ±SE. Asterisks represent the statistical significance between treatments as judged by Student’s *t*-test: **P < 0.001 and ***P < 0.0001.

To confirm the radioimaging results, we also performed direct scintillation counting experiment using individual roots. A noticeable increase in ^74,73^As activity was observed in *pin2*/*eir1-1* roots for all tested arsenite concentrations (Figure 4C). Collectively, these results suggest the possibility that As(III) species could serve as a transport substrate for PIN2 and hence loss of PIN2 functioning would result in higher intracellular accumulation of arsenite.

### Arsenic content is higher in *pin2*/*eir1-1* mutant

Due to the low specific activity of ^74,73^As, the radioactive transport assay was conducted on whole plant roots. However, PIN2 is preferentially expressed in meristem and elongation zone (Xu and Scheres, 2005; Shibasaki et al., 2009). To confirm the functional role of PIN2 in As(III) transport, we determined arsenic concentrations in the 5mm-long root tips after short term (2h) exogenous arsenite treatment using ICP-MS. Compared with wild-type, almost two-fold increase in arsenic accumulation was observed in *pin2/eir1-1* mutant plants (Figure 4D). Similar results were observed for a long term (3d) arsenite treatment (Supplemental Figure 9).

### PIN2 is incapable of transporting arsenite in *ycf1*Δ *acr3*Δ deletion mutant of *S. cerevisiae*

Heterologous expression system is another approach to assess the protein transporter activity. Besides the bacterial arsenite transporters ArsB, AtPINs show homology (18%) to arsenite transporters Acr3 from *S. cerevisiae* (Table 1). Arsenite export by Acr3p is one of the most important arsenic detoxification mechanisms discovered in *S. cerevisiae* (Ghosh et al., 2002). Another protein affecting arsenite resistance of yeast, Ycf1p, is located at the vacuolar membrane. Ycf1p, a member of the multidrug resistance (MRP) group of the ABC superfamily of drug resistance ATPases, mediates the active transport of glutathione-conjugated toxic compounds, including the product of arsenite sequestration, As(GS)_3_ in the yeast vacuole (Ghosh et al., 2002).

Hence, for testing the arsenite transport activity of PIN2, we selected the yeast strain lacking both Acr3p and Ycf1p (*ycf1*Δ *acr3*Δ). Expression of Acr3 in *ycf1*Δ *acr3*Δ did result in increased resistance to arsenite in growth assay and reduced accumulation of arsenite in transport assay. However, PIN2 did not show any arsenite transport activity (Supplemental Figure 10). These results suggest that even if PIN2 may be involved in arsenite transport in plants, it is not functional as such in *S. cerevisiae*. This finding is not inconsistent as in numerous studies it has been shown that the expression of plant proteins in heterologous system widely varies depending on the system that is used, and in many cases plant proteins either do not express in heterologous system or not showing the same functionality (Dreher et al., 2006; Ma et al., 2008; Barbosa et al., 2018). However, the members of PINs have been expressed in a variety of heterologous systems, including *S. cerevisiae*, fission yeast *Shizosaccharomyces pombe*, human *HeLa* cultured cells, and *Xenopus laevis* oocytes, as reviewed in (Barbosa et al., 2018). Arabidopsis PIN1, PIN2 and PIN7 were previously successfully expressed in *S. pombe* and showed comparable expression levels and IAA export activity (Yang and Murphy, 2009). Unfortunately, the proteins from the Acr3 family of transporters are widely distributed in prokaryotes and fungi with the exception of *S. pombe* (Wysocki et al., 2003; Mansour et al., 2007), and hence the current assay protocol need to be modified to be used for studies of arsenite using this system. Testing of the available *S. pombe* strains and other heterologous systems to study properties of PIN2 as transporter of trivalent arsenicals will be a topic of our future research.

### Speciation and localization of arsenic by Synchrotron X-ray Fluorescence Imaging

High resolution synchrotron X-ray fluorescence imaging (XFI) coupled with XAS is a powerful technique to identify the localization and chemical speciation of metals and metalloids in situ (Pickering et al., 2000). In this study, application of the synchrotron techniques pursued the following goals: 1) to compare the patterns of arsenic distribution and its relative concentrations in the roots of arsenite-exposed wild type and *pin2*/*eir1-1* mutant plants, and 2) to determine chemical speciation of arsenic accumulated in the different tissues of wild type and mutant plants, using micro-XAS and bulk XAS techniques.

Since the short term and long term treatments with exogenous arsenite essentially produced similar trends in PIN2 trafficking (Figure 3 and Supplemental Figure 8), and arsenic accumulation (Figure 4D and Supplemental Figure 9), in the synchrotron experiments we used long term (3 day) arsenite-treated plants to simplify plant transportation.

XFI imaging revealed a striking difference in arsenite localization in wild-type and *pin2/eir1-1* roots (Figure 5A and 5B). While arsenic accumulated at the very end of the root tip of arsenite exposed *pin2/eir1-1* mutant, the arsenic distribution seems to be more diffuse in the root apical meristem of wild-type (Figure 5A and 5B). Calculations of arsenic areal densities in the comparable portions of the apical root meristem in arsenite-exposed wild type and *pin2/eir1-1* root samples using XFI maps revealed 2-3 times higher mean values for arsenic in *pin2/eir1-1* root tips compared with that of wild-type (Table 2 and Figure 5F). These results are consistent with the observed difference in accumulation of arsenic in wild type and *pin2/eir1-1* root tips and whole root determined by ICP-MS (Figure 4D and Supplemental Figure 9), confirming that mutations in PIN2 lead to altered arsenic response in *A. thaliana*.

**Figure 5.**
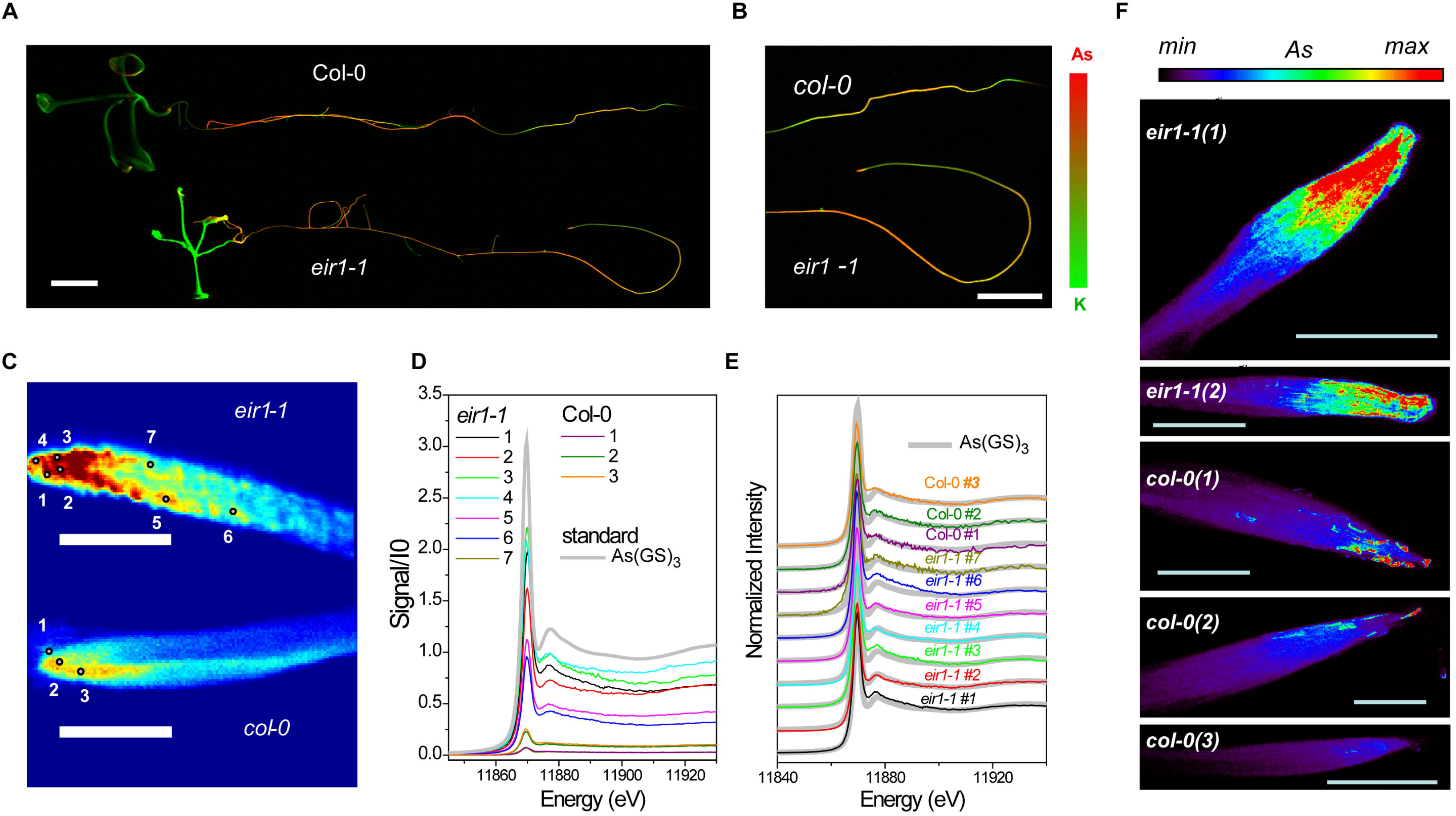
As accumulation and distribution in roots of *Arabidopsis thaliana* exposed to 10 μM arsenite are influenced by PIN2. **(A)** Combined XFI As and potassium (K) elemental distributions in whole Arabidopsis plants. As is denoted by red and potassium by green with brighter colors corresponding higher concentrations. As (and K) intensities are in a common scale for two specimens. The samples were scanned with 35 μm step at the beamline 10-2 (SSRL). Spatial scale bar represents 3.5 mm. **(B)** The images of roots shown in **(A)** are magnified by 1.4 times to show differences in the As distribution in the apical part in the root meristem in Col-0 and *eir1-1*. **(C)** The high resolution As XFI maps of the apical root meristem demonstrate higher accumulation of As in *eir1-1* mutant and less concentrated and more diffused distribution in Col-0. Arsenic intensity is in common scale for two specimens. Brighter colors correspond to higher concentrations. The circular markers denote spatial points that were selected for collection of As micro-XAS in roots. The samples were scanned with 2 um step at the beamline 2-3 (SSRL). (Spatial scale bar – 100 um). **(D)** Micro-XAS As-K near-edge spectra collected at the points of the apical root meristem in Col-0 and *eir1-1* roots as marked in (C). The spectra are normalized by intensity of the incident radiation but otherwise are not processed. Apart from *eir1-1 #7*, micro-XAS As-K spectra in *eir1-1* are much more intense compared with Col-0. **(E)** Same near-edge XAS As-K spectra as in **(D)**, collected in the root points shown in (C), with background removed, normalized by the intensity of the incident radiation and the edge jump. All these spectra show a high similarity to As III - thiolated species, best represented by As(GS)_3_ standard. **(F)** High resolution XFI As areal density distributions in hydrated root specimens of Arabidopsis collected at the beamline 2-ID-E (APS) with spatial resolution 1 μm × 1 μm.

**Table 2:** Areal densities (in ng/cm^2^) As, Fe, Zn, Ca.

| element | As |  | Fe |  | Zn |  | Ca |  |
| --- | --- | --- | --- | --- | --- | --- | --- | --- |
| plant/zone | mean | Median | mean | median | mean | median | Mean | median |
| **Col-0 + 10 μM arsenite** |  |  |  |  |  |  |  |  |
| Z1 | 1750 | 1163 | 1197 | 885 | 489 | 517 | 13418 | 13894 |
| Z2 | 1068 | 730 | 873 | 720 | 543 | 565 | 13142 | 13660 |
| **Col-0 + 10 μM arsenite** |  |  |  |  |  |  |  |  |
| Z1 | 856 | 835 | 1424 | 1394 | 665 | 655 | 12215 | 11679 |
| Z2 | 535 | 483 | 1269 | 1272 | 638 | 642 | 9882 | 9716 |
| **Col-0 + 10 μM arsenite** |  |  |  |  |  |  |  |  |
| Z1 | 1208 | 1134 | 634 | 621 | 509 | 505 | 10640 | 10047 |
| Z2 | 940 | 838 | 672 | 654 | 678 | 691 | 13091 | 12782 |
| ***eir1-1* + 10 μM arsenite** |  |  |  |  |  |  |  |  |
| Z1 | 5252 | 5354 | 1714 | 1539 | 815 | 845 | 11291 | 11078 |
| Z2 | 4927 | 4730 | 892 | 638 | 1199 | 1200 | 12693 | 12593 |
| ***eir1-1* + 10 μM arsenite** |  |  |  |  |  |  |  |  |
| Z1 | 3093 | 2953 | 901 | 836 | 619 | 602 | 11090 | 11705 |
| Z2 | 3460 | 3368 | 986 | 899 | 587 | 552 | 10843 | 10861 |
| **Col -control** |  |  |  |  |  |  |  |  |
| Z1 | 7 | 0 | 1272 | 1161 | 504 | 501 | 11925 | 11730 |
| Z2 | 7 | 1 | 1331 | 1274 | 775 | 737 | 18797 | 18409 |
| ***eir1-1* control** |  |  |  |  |  |  |  |  |
| Z1 | 6 | 0 | 1189 | 971 | 265 | 223 | 7701 | 6545 |
| Z2 | 4 | 0 | 764 | 640 | 219 | 187 | 4609 | 3800 |

The areal density was calculated from the elemental maps obtained from 2-ID-E images of col-0
and *eir1-1* in presence or absence of 10 μM arsenite.
Z1 – denotes the root apical meristem.
Z2- denotes the transition zone.

The micro-XAS analysis of As K-edge (near edge spectra) was conducted at various parts of the roots as shown in Figure 5C. The results of principal component analysis (PCA) and least-square fitting of linear components of near-edge micro-XAS show that the majority of the arsenic is best represented by As(III)-tris-thiolate complex (Figure 5D and 5E). For arsenic imaging, a similar thiolate complex was observed in Indian Mustard (*Brassica juncea*), which belongs to the same *Brassicaceae* family as *A. thaliana* (Pickering et al., 2000).

The XFI elemental maps at the micron and sub-micron scale allowed us to compare accumulation of arsenic and other biologically important elements in the root apical meristem of *pin2*/*eir1-1* and wild type. The elemental analysis data are presented in Table 2. Root tips of arsenite-exposed *pin2/eir1-1* showed 2-3 fold higher arsenic accumulation compared with the wild-type root tip. For other elements like Fe, Zn and Ca, no large differences in accumulation were observed between wild-type and *pin2/eir1-1* (Table 2 and Supplemental Figure 11, 12). Collectively, these results support the results of the radioactive arsenite transport assay and ICP-MS measurements that arsenic accumulates in higher levels in root meristem zone of *pin2/eir1-1* as compared to wild-type plants.

## Discussion

In this work, we provide a new insight into the role of auxin efflux carrier PIN2 in regulating root arsenite response as well as in facilitating the intracellular transport of As (III) species. Several lines of molecular and cellular evidence suggest that response of *Arabidopsis* root to arsenite but not arsenate is tightly linked to altered intracellular auxin homeostasis, regulated by auxin efflux carrier PIN2. Consistently, the loss-of-function mutant *pin2* plants exhibit striking phenotypic changes in the root morphology, and accumulated 2-3 fold higher arsenic concentrations in root apices compared to that of wild-type plants. The arsenite response in root was found to be linked to altered auxin homeostasis. Arsenite inhibited the shootward auxin transport and subsequently the intracellular auxin distribution, which was supported by the observed altered signal intensities and distribution patterns in auxin marker lines. We further demonstrated that arsenited-induced change in auxin distribution is directly linked to the intracellular trafficking of PIN2. Analysis of cellular localization and trafficking of PIN2 and two other membrane proteins PIN1 and LTI6b, all of which is trafficked from the plasma membrane to endosome through a BFA regulated pathway (Geldner et al., 2001; Kurup et al., 2005; Shibasaki et al., 2009), revealed that arsenite specifically inhibits the PIN2 trafficking as in presence of Arsenite, formation of BFA bodies was abolished only for PIN2 but not for PIN1 and LTI6b.

Comparison of arsenic transport dynamics and arsenic accumulation in wild-type and *pin2/eir1-1* mutant plants by *in planta* transport assay, ICP-MS and high resolution synchrotron fluorescence imaging coupled with micro-XAS at selected root meristem provides evidence that arsenite efflux in *A. thaliana* is linked to PIN2 functioning. It is important to note that the in planta transport assay and ICP-MS analyses were performed for a brief period (2h), which rule out the possibility of any long term indirect effect of PIN2 mutation in regulating the transport of arsenite. The highest As accumulation areas in the elemental maps of *pin2/eir1-1* mutants obtained by synchrotron XFI were in the root cap and epidermis of apical meristem, the same tissues where PIN2 would be normally expressed in wild type. Moreover, only the arsenic content but not the other elements such as Fe, Zn and Ca content is altered in *pin2/eir1-1* root tip and the epidermis/cortical zones of apical meristem compared to wild-type. This specific increased accumulation of arsenite in the root tip clearly rules out the possibility of any pleiotropic phenotype of PIN2 mutation affecting the movement/sensitivity of other inorganic ions indirectly and demonstrates the importance of PIN2 in cellular arsenite efflux.

In contrary to an earlier study (Krishnamurthy and Rathinasabapathi, 2013), where it was claimed that both *pin1* and *pin2* showed hypersensitive response to arsenite-induced root growth inhibition, we did not find any effect of arsenite on PIN1 neither in the root growth nor in trafficking assays. We found several other discrepancies in this work. For instance, the author claimed that exogenous IAA treatment alleviates arsenite tolerance in *aux1*, which is not explainable as numerous studies showed that *aux1* is IAA resistant and IAA uptake is significantly reduced in *aux1* (Pickett et al., 1990; Marchant et al., 1999; Rahman et al., 2001). The authors also claimed that arsenite inhibits auxin uptake. However, the authors performed acropetal and basipetal transport experiments which are completely different from the auxin uptake experiment and do not truly represent the auxin uptake status of the root (Rashotte et al., 2000; Rashotte et al., 2001; Lewis et al., 2007; Shibasaki et al., 2009; Hanzawa et al., 2013). The only substantial difference in these two works was the plant growth condition; while they used an alternating light/dark regime, we used continuous light. Hence, the results presented in (Krishnamurthy and Rathinasabapathi, 2013) work should be carefully inerpreted.

It might seem paradoxical that arsenite specifically targets the intracellular cycling of PIN2, although PIN2 and PIN1 show similar homology to several arsenite transporters Sc ACR3, PV ACR3 and arsB. However, this is not inconsistent as PIN1 and PIN2 use distinct pathways for trafficking and cellular targeting (Krecek et al., 2009). For instance, in roots of *A. thaliana*, PIN1 is expressed only in the central cell files, where it always shows a polarization towards the rootward domain of the plasma membrane (Geldner et al., 2001). On the other hand, PIN2 is expressed in the lateral root cap cells, epidermis and cortex with a mixed polarity. In LRC, epidermis and mature cortical cells, PIN2 shows a polarization towards the shootward domain of plasma membrane as opposed to PIN1 polarization, while in meristematic cortical cells, it shows rootward polarization like PIN1 (Rahman et al., 2007; Rahman et al., 2010). The subcellular targeting mechanisms of PIN1 and PIN2 are also distinct. Newly synthesized nonpolar PIN1 and meristematic cortical PIN2 achieve the rootward polarity through ARF-GEF, such as GNOM, and the phosphorylation status of the protein, which is regulated by the counter balancing activities of PINOID kinase and protein phosphatase 2A. Rootward polarity of PIN1 and cortical PIN2 can be reversed to shootward by altering the phosphorylation status of the protein through over expression of PID kinase or by reducing the protein phosphatase activity through genetic or pharmacological approaches (Friml et al., 2004; Michniewicz et al., 2007; Kleine-Vehn and Friml, 2008; Rahman et al., 2010). However, PIN1 and meristematic cortical PIN2 showed differential phosporylation requirements for relocalization towards shootward domain (Rahman et al., 2010). Moreover, polarization of PIN2 in LRC, and in epidermal cells is completely independent of this pathway (Friml et al., 2004; Rahman et al., 2010; Hanzawa et al., 2013). These results exclusively suggest that trafficking pathways of PIN1 and PIN2 are distinct and support our observation that arsenite selectively targets the machinery that only regulates PIN2 trafficking.

In many species, removal of arsenite from the cytoplasm is mainly regulated by energy coupled systems. In bacteria and yeast, ATP coupled ArsAB, and H^+^ coupled ArsB and Acr3 function in active extrusion of arsenite (Yang et al., 2012). ArsB can also function as a subunit of the ArsAB As(III)-translocating ATPase, an ATP-driven efflux pump. In this complex, As(III) binding to three thiols of ArsA induces a conformational change that increases the rate of ATP hydrolysis and, consequently, the rate of As(III) extrusion by the ArsAB pump. Experimental evidence in oocytes suggests that efflux of silicon by Lsi2 is an energy-dependent active process driven by the proton gradient. However, expression of Lsi2 in oocytes, yeast and bacteria did not show any arsenite transport activity (Ma et al., 2008). Similarly, expression of PIN2 in *S.cerevisea* did not show any arsenite transport activity either. These results also highlight the possibility that these proteins function *in planta* is aided by other proteins, which are absent in the heterologous system and hence failed to show the expected transport activity.

PINs have been shown to interact with another group of membrane transporter family protein PGP/ABCBs (ATP-binding cassette transporters of the B subfamily), involved in auxin transport (Blakeslee et al., 2007; Zazímalová et al., 2010; Cho et al., 2012; Geisler et al., 2017). “Concerted” interactions have been shown between ABCB19 and PIN1 or ABCB1/ABCB4 and PIN2 in auxin transport in polar PM domains, where ABCBs and PINs can physically interact (Blakeslee et al., 2007; Titapiwatanakun et al., 2009; Cho et al., 2012). It was also proposed that while members of the ABCB and PIN families can function as independent auxin transport catalysts, “a strict co-operative or mutual functionality” cannot be excluded (Geisler et al., 2017). In many organisms, proteins from PGP/ABCB/MDR group are involved in active efflux of various xenobiotics, including metals, from the cells. In fact, MDR1/ABCB1 gene codes a P-glycoprotein was the first ABC transporter correlated with arsenic resistance in human renal carcinoma cells (Maciaszczyk-Dziubinska et al., 2012). We speculate that selected plant ABCB transporters could also interact with PIN2 in a manner analogous to ArsAB ATPase complex in extrusion of arsenic.

Although the uptake mechanism of arsenite in plant is largely understood, efflux mechanisms as well as root response mechanism to arsenite remain elusive. The findings of the present study that PIN2 and auxin are intrinsically involved in regulating arsenite responses in roots, and that PIN2 functions as a possible arsenite efflux transporter open a new door to our understanding. Future studies aiming to identify the substrates of arsenite that are effluxed by PIN2 or Lsi2 through structural NMR study, and elucidating the substrate binding domains in these proteins would further enhance our understanding of arsenite efflux mechanism.

## Materials and Methods

### Plant materials

All lines except, *pin1-3* [Ler background (Bennett et al., 1995)], EGPP-LTI6b [C24 background (Kurup et al., 2005)], are in the Columbia background of *Arabidopsis thaliana* (L.). *eir1-4, eir1-4* PIN2::PIN2 and eir1-4 35S::PIN2 were described earlier (Abas et al., 2006, Retzer et al., 2017). *PIN2–GFP* (Xu and Scheres, 2005) was the gift of B. Scheres (University of Utrecht, The Netherlands); DII-VENUS (Brunoud et al., 2012) was a gift from Malcolm Bennett (University of Nottingham, UK). *pin1-3*, *pin3-4*, *pin4-3* and GFP-LTI6b were provided by Gloria Muday (Wake Forest University, Winsto-Salem, NC, USA). Col-0, *eir1-1* and PIN1-GFP were obtained from the Arabidopsis Biological Resource Centre (Columbus, OH, USA).

### Growth conditions

Surface-sterilized seeds were germinated and grown for 5 days containing 1% w/v sucrose and 1% w/v agar (Difco Bacto agar, BD laboratories; http://www.bd.com) in a growth chamber (NK system, LH-70CCFL-CT, Japan) at 23° C under continuous white light (at an irradiance of 80-100 μmol m^−2^ s^−1^; Shibasaki et al., 2009). The seedlings were grown vertically for 5 days and then transferred to nesw plates with or without arsenite and arsenate, and incubated for various time lengths under continuous light at irradiance of 80-100 μmol m^−2^ s^−1^ (NK system, LH-1-120.S, Japan). After the incubation, the pictures of the seedlings were taken using a digital camera (Canon; Power Shot A 640, http://canon.jp/) and root elongation of the seedlings were analyzed by an image analyzing software Image J (http://rsb.info.nih.gov/ij/). *pin1-3* was maintained as heterozygous and homozygous seedlings were selected using the fused cotyledon phenotype as described earlier (Aida et al., 2002).

### Chemicals

Sodium (meta) arsenite (NaAsO_2_) and sodium arsenate dibasic heptahydrate (Na_2_HAsO_4_^.^7H O) were purchased from Kanto Chemical Co. (Tokyo, Japan). BFA was purchased from Sigma-Aldrich Chemical Co.(USA). [^3^H] IAA (20 Ci mmol^−1^) was purchased from American Radiolabeled Chemicals, Inc. (St. Louis, MO, U.S.A., http://www.arc-inc.com). The chemicals for growth media and standards used in synchrotron experiments were purchased from Sigma-Aldrich Chemical Co. (Canada). Other chemicals were from Wako Pure Chemical Industries (http://www.wako-chem.co.jp/).

### Bioinformatics analysis

Protein sequences of AtPIN1 (AT1G73590), AtPIN2 (AT5G57090), AtPIN3 (AT1G70940), AtPIN4 (AT2G01420) and AtPIN7 (AT1G23080) were collected from TAIR (www.arabidopsis.org). *Saccharomyces cerevisiae* Acr3 (190409919), *Pteris vittata* Acr3 (310768536), *Escherichia coli* arsenite transporter ArsB (1703365), and *Oryza sativa* silicon transporter Lsi2 (296936086) were collected from NCBI (www.ncbi.nlm.nih.gov) protein database. Multiple sequence alignment (Supplemental Figure 2 and 3) and identity matrix (Table 1; Supplemental Tables 1, 2 and 3) are generated using Clustal Omega (Sievers et al., 2011) (http://www.ebi.ac.uk/Tools/msa/clustalo/) multiple sequence alignment tool. Cladogram of plasma membrane residing PIN proteins (AtPIN1, AtPIN2, AtPIN3, AtPIN4, AtPIN7) (Supplemental Figure 1) from *Arabidopsis thaliana* were constructed using MEGA6 (Tamura et al., 2013) (Molecular Evolutionary Genetics Analysis) software based on Neighbor Joining method and 1000 bootstrap test.

### Gravitropism assay

Root tip reorientation was assayed as described earlier (Rahman et al., 2010). In brief, 5-day-old vertically grown seedlings were transferred to new square plates in presence or absence of arsenite. After the transfer, the roots were gravistimulated at 23°C by rotating the plate 90°. To measure the curvature of roots and elongation, photographs of plates were taken at specific time points after reorientation using a digital camera (Canon; Power Shot A 640, http://canon.jp/) and analyzed by an image analyzing software Image J (http://rsb.info.nih.gov/ij/). Data were obtained from three biological replicates.

### Transport assay

#### i) Auxin transport assay

5-day-old vertically grown Arabidopsis seedlings were transferred to agar plate and incubated with or without 10μM arsenite for 3 days. Shootward auxin transport was measured as described earlier (Shibasaki et al., 2009). In brief, a donor drop was prepared by mixing 0.5 μM [^3^H] IAA (3.7 MBq ml^−1^) in 1.5% agar containing MES buffer solution. The donor drop was placed on the edge of the root tip. Plates were then incubated vertically at 23°C for 2 h. To measure auxin transport, 5 mm root segments away from the apical 2 mm were carefully cut and soaked for overnight in 4 mL liquid scintillation fluid (ULTIMA GOLD, PerkinElmer, USA), and the radioactivity was measured with a scintillation counter (model LS6500, Beckman ACSII, USA Instruments, Fullerton, CA). Data were obtained from at least three biological replicates.

#### ii) Arsenite transport assay

^74,73^As (III) prepared from radioactive arsenic (^74,73^As) was used for transport assay. (^74,73^As (III) preparation method is described in detail in supplemental method section). 5-day-old vertically grown Col-0 and *eir1-1* seedlings were incubated in Hoagland solution containing 0.1 and 10 μM labeled ^74,73^As (III) (approximately 5KBq ml^−1^) for 2h and transferred to a fresh plate to separate root and shoot samples. For radioluminography, the seedling roots were incubated in 0.1 and 10 μM and then placed on a Kraft paper using double-sided tape. The sample was exposed to an imaging plate (IP; BASIP MS, GE Healthcare Lifescience) and the radiation distribution was visualized by FLA-5000 Image Analyzer (FUJIFILM). Arsenic contents in seedlings were calculated from the value of photostimulated luminescence in the imaging data. Data were obtained from three biological replicates.

For liquid scintillation count, individual whole root sample was taken in separate tubes and 10 individual roots were considered for each treatment. Individual root sample was collected in a vial with scintillation cocktail (MICROSCINT 40, PerkinElmer) and ^74,73^As activity was measured by liquid scintillation counter (Tri-Carb 4810 TR, PerkinElmer) with the window of 0 to 2000 keV.

### Live cell imaging and GUS staining

For live cell microscopy, five-day-old GFP or DII-VENUS transgenic seedlings were used. For short term BFA treatment, five-day-old seedlings were incubated in 10 μM arsenite for 2h and then subjected to 20 μM BFA for 40 minutes. For long term five-day-old seedlings were incubated with or without 10 μM arsenite for additional 3 days. After the incubation, BFA treatment was performed as described above. After the BFA incubation, the seedlings were mounted in liquid growth medium on a cover glass for observation on a Nikon laser scanning microscope (Eclipse Ti equipped with Nikon C2 Si laser scanning unit) and imaged with as x40 water immersion objective.

The accumulation of BFA bodies in PIN2-GFP was quantified in the transition area of the root tip as described earlier (Shibasaki et al., 2009). The pictures were taken approximately at the same place of the root and using same confocal settings. The number of BFA bodies and cells were counted for each root and expressed as BFA body/cell. Data were obtained from at least three biological replicates. DII-Venus imaging and GUS staining are described in supplementary section.

### Measurement of arsenic content in plants

For short term treatment, five day old wild-type and *pin2/eir1-1* seedlings were transferred to Hoagland solution containing 10 μM and 100 μM arsenite and incubated for 2h. After incubation, roots were washed three times and 5 mm root tip were cut dried and weighed. 20 root tips were collected into a vial. After digestion by nitric acid, As content in the sample solution was measured by inductively coupled plasma mass spectrometry (ICP-MS) (NexION 350S, PerkinElmer). For the long term treatment, 5 day old seedlings were treated for 3 days in arsenite-spiked agarose growth media and As content in roots and shoots was measured as described above.

### Expression of PIN2 in *ycf1*Δ *acr3*Δ mutant of *S. cerevisiae*

Since arsenite resistance in *S. cerevisiae* is largely mediated by Acr3p and Ycf1p, we used the yeast strain lacking both Ycf1p and Acr3p (*ycf1*Δ *acr3*Δ). PIN2 and Acr3p were cloned in pKT10 vector, transformed into the double knock out strain and selected with AHCW/Glc medium. Preparation of double knock out strain, cloning of PIN2 and Acr3p, growth and transport assays are described in detail in supplementary method.

### X-ray fluorescence imaging (XFI) and X-ray absorption spectroscopy (XAS)

Synchrotron XFI was performed at the Stanford Synchrotron Radiation Lightsource (SSRL), Stanford University, Menlo Park, CA, USA (beamlines 10-2 and 2-3) and at the Advanced Photon Source (APS), Argonne National Laboratory, Lemont, IL, USA (beamline 2-ID-E). Micro-XAS analysis of selected arsenic hot spots in the samples at a room temperature was performed at the beamline 2-3 (SSRL), and bulk As K-edge XAS of frozen plant tissues and growth medium was conducted in a helium cryostat environment at the beamline 7-3 (SSRL).

### Preparation of Samples for synchrotron XFI imaging

*Col-0* and *pin2/eir1-1* were grown as described above. 5-d-old seedlings were treated with 10μM arsenite in growth medium for 3 days. After the treatment, control and arsenite-exposed plants were harvested, washed in de-ionized (DW) water and either flash frozen for consecutive bulk XAS analysis or imaged as described below and, in more detail, in Supplemental information.

### XFI of whole plants

Elemental mapping of whole plants using XFI was performed at the wiggler beamline 10-2 (SSRL). The samples were mounted at 45° to the incident beam and raster scanned (pixel size 35 μm × 35 μm) with beam dwell time 100 ms per pixel (Supplemental Figure 13, see Supplemental information for details). Fluorescent energy windows were centered for As, Fe and other elements of interest (P, S, K, Ca, Zn, Cu, Mn).

### Micro XFI and Micro-XAS of selected plant tissues

Higher resolution imaging (2 – 10 μm) of selected tissues of interest (areas of the apical root meristem of live main roots) were performed at the bending magnet beamline 2-3 (SSRL) with the same geometric and fluorescent energy detection setup as described for the beamline 10-2 above. After high resolution images were obtained, we collected micro-XAS As-K near-edge spectra in the As ‘hotspots’ and other areas of interest.

### Bulk XAS at the beamline 7-3 (SSRL)

The pre-washed roots and freshly made agar medium were flash-frozen in the liquid nitrogen in separate Lucite cells and measured at the wiggler beamline 7-3 (SSRL) at ~ 10K using helium cryostat. The micro-XAS and bulk XAS spectra were analyzed using EXAFSPAK (http://ssrl.slac.stanford.edu/~george/exafspak/manual.pdf) suite of programs (see Supplemental information for the details).

### X-ray Fluorescence Microscopy at the APS (2-ID-E station)

X-ray fluorescence microscopy (XFM) of fresh hydrated root apical meristem with 1 micron and 300 nm resolution was conducted at the 2-ID-E X-ray fluorescence microprobe station of the Advanced Photon Source (APS), as described in SI. The fitting of two‐dimensional elemental maps and analysis of the regions of interest (ROI) analysis were performed using MAPS software (Nietzold et al., 2018).

### Statistical Analysis

Results are expressed as the means ± S.E.M from appropriate number of experiments as described in the figure legends. Two-tailed Student t-test was used to analyze statistical significance.

## Author contributions

A.R., O.P., G.N.G., I.J.P., C.L., designed the research. M.A.A., M.S.,M.A., C.D.K., O.P., N.V.D., S.N., O.A., T.K., and A.R. performed the experiments. K.T., K.M., and K.N., produced the radioactive arsenite. M.A.A., O.A., O.P., K.T.,and A.R., analyzed the data. M.A.A, O.P, N.V.D., G.N.G, I.J.P, K.T., T.K., and A.R., wrote the paper.

## Acknowledgement

This work was supported in part by the Iwate University President Fund (to A.R.), and Global Innovation Fund, University of Saskatchewan (to I.J.P., G.N.G., A.R). Research at the University of Saskatchewan is supported by grants from the Natural Sciences and Engineering Research Council of Canada (G.N.G., I.J.P.), the Saskatchewan Health Research Foundation (G.N.G., I.J.P.), The University of Saskatchewan, and by Canada Research Chairs (G.N.G., I.J.P.).

Resource of the Advanced Photon Source is supported by the U.S. Department of Energy (DOE) Office of Science User Facility operated for the DOE Office of Science by Argonne National Laboratory under Contract No. DE-AC02-06CH11357.

Use of the Stanford Synchrotron Radiation Lightsource, SLAC National Accelerator Laboratory, is supported by the U.S. Department of Energy, Office of Science, Office of Basic Energy Sciences under Contract No. DE-AC02-76SF00515. The SSRL Structural Molecular Biology Program is supported by the DOE Office of Biological and Environmental Research, and by the National Institutes of Health, National Institute of General Medical Sciences (including P41GM103393). The contents of this publication are solely the responsibility of the authors and do not necessarily represent the official views of NIGMS or NIH.

