## Supplemental Information for "PIN FORMED 2 facilitates the transport of Arsenite in *Arabidopsis thaliana*"

**Methods:**

**GUS staining**

GUS staining was performed as described earlier (Okamoto et al., 2008). In brief, five-day-old seedlings were transferred to Hoagland solution in presence or absence of arsenite and incubated for 1h and 2h respectively. After the incubation, seedlings were transferred to GUS staining buffer (100mM sodium phosphate, pH7.0, 10 mM EDTA, 0.5 mM potassium ferricyanide, 0.5 mM potassium ferrocyanide, 0.1% Triton X-100) containing 1 mM X-gluc and incubated at 37°C in the dark for 1 h. The roots were imaged with a light microscope (Nikon, Diaphot, Japan, www.nikon.co.jp) equipped with a digital camera control unit (DIGITAL SIGHT (DS-L2); Nikon, Japan, www.nikon.co.jp).

**Live cell Imaging**

Five-day-old DII-Venus seedlings were treated with 10mM Arsenite for 3d. After the incubation, fluorescence intensities were measured as described earlier (Hanzawa et al., 2013) by drawing a region of interest (ROI) in the images obtained from the live-cell imaging using Image J software.

**Gene Expression Analysis**

5-day-old vertically grown Arabidopsis seedlings were transferred to new agar plate and grown vertically in presence or absence of arsenite for 3 days under continuous light. After the incubation, RNA was extracted from the root tissue using PlantRNA extraction kit (Qiagen, USA) and tested for quality and quantity. Each RNA concentration was normalized with RNase free water. 1μg RNA was applied to synthesize single strand cDNA using Rever Tra Ace qPCR RT master mix (Toyobo, Japan). Quantitative PCR reactions were performed using the Takara TP-850 thermal cycler (Takara Bio, Japan) and SYBR® Premix Ex Taq qPCR kit (Takara Bio, Japan).The reaction was performed as per manufacturer's instruction. For quantification of PINs expression, we used the 2-ΔΔCT (cycle threshold) method (Livak et al., 2001). Data were obtained from three biological replicates.

Primer sequences used to analyze *PIN* expression are as follows:

*PIN1* Left primer ATCTTCACATGTTTGTGTGG
Right Primer TCGTCTTTGTTACCGAAACT
*PIN2* Left primer AGATGCCAACGATAATGAGT
Right Primer AGTAATCACCTGAACGATGG

**Preparation of radioactive arsenite**

Radio-arsenic (74,73As) was produced by irradiating an elemental natGa with alpha-particles (on-target). Radio-arsenic was extracted by a cocktail of HCL and hydrogen peroxide. After purification, the radio-arsenic was reduced to arsenite by titanium trichloride. The radioactive isotopes found in the product was 71As, 72As 73As and 74As. Because the half-lives of 71As (65 h), 72As (26 h) are short enough to decay out after several weeks, the radio-arsenic consisted of 74,73As in the present study.

After reducing arsenate (V) into arsenite (III), TLC assay was performed to confirm the chemical form of As isotope by the modified method of Resano et al. (2007). In short, 2 μl of As isotope solution was applied on the TLC plate (TLC PEI cellulose F., Merck Millipore). After drying, the TLC plate was developed with a mixture of acetone : acetic acid : water (2:2:1 v/v/v). As isotope was detected by radioluminography using BAS IP MS (FUJIFILM). We confirmed that >99% of total As isotope was arsenite (III).

**Arsenite transport assay in *Saccharomyces cerevisiae***

In *Saccharomyces cerevisiae*, two proteins are involved in As(III) transport: Acr3p and Ycf1p (Wysockiet al., 1997; Ghosh et al., 1999). Therefore, to measure the transport activity, we used yeast strain lacking both Ycf1p and Acr3p (*ycf1Δ acr3Δ*).

The yeast mutant was prepared as follows. To disrupt Ycf1, pAG32 (Goldstein and McCusker, 1999) was amplified by PCR using 5’CATGTGGGTTGGCGTGATTATACTAGTTATTATGATGCCAAGCTGAAGCTTCGTACGCTG-3’ and 5’CCAGATAAGGAGATCCCTTTCTCGCCAACTAATGTCTTATCTCCATAGGCCACTAGTGGATCTG-3’. The amplified fragment was transformed into *ycf1Δ* and selected aureobasidin

A (Takara) resistant colony, which is *ycf1Δ acr3Δ.*

PIN2 cDNA was amplified with primers

5'-ACGAATTCATGATCACCGGCAAAGACATG-3' and

5'-ACGTCGACTTAAAGCCCCAAAAGAACGTAG-3'.

As a positive control ACR3 gene was amplified from the gDNA of yeast strain BY4741 with primers 5'-GCGAATTCATGTCAGAAGATCAAAAAAG -3'

and 5'-GCGTCGACTTAATTTCTATTGTTCCATA -3' .

The resulting PCR fragments were digested with *Eco*RI and *Sal*I, and inserted into pKT10 (Tanaka et al. 1990) at the *Eco*RI–*Sal*I site. The final vector construct was confirmed by sequencing before transforming into yeast for transport experiment.

For the measurement of As (III) transport, the plasmids were transformed into double mutant and selected with AHCW/Glc medium (Ueoka-Nakanishi, 1999). The As (III) uptake assay was performed as follows. The yeast transformants were incubated with AHCW/Glc overnight and then precultured with SD medium (0.67% yeast nitrogen base w/o amino acid, 2% glucose) supplemented with auxotrophic requirements until $OD_{600}$ reached about 0.5. The yeast were collected by centrifugation and suspended in SD medium. 3-4 x $10^7$ cells were incubated with SD medium containing 100 μM As (III) with shaking. After 1 h incubation, cells were collected with centrifugation and washed with ice-cold SD medium three times. The yeasts were dried at 65˚C and digested with $HNO_3$. After the digestion, the samples were dissolved in 0.08 N $HNO_3$ containing 2 ppb indium (In) and applied for ICP-MS (Agilent 7800).

**Synchrotron X-ray Fluorescence Imaging (XFI) and X-ray Absorption Spectroscopy (XAS)**

**XFI of whole plants**

Elemental mapping of whole plants using XFI was performed at the wiggler beamline 10-2 (SSRL) using a Si(111) double-crystal monochromator (phi =90°) with a Rh-coated mirror for beam focusing. Plants were carefully removed from agar growth medium, washed with the deionized water and gently patted with Kimtech® wipes. The whole plants were spread on the surface of a metal-free PET plastic sheet (thickness 0.175 mm) and covered with stretched 0.16 mil (4 μm) thick Ultralene ® XRF film which was fixed with metal-free adhesive tape along the perimeter of the sample.

The samples were mounted at 45° to the incident beam (13450 eV) which was focused through an aperture upstream of the I0 ion chamber, producing the beam spot 50 μm x 50 μm size.

X-ray fluorescence was detected using a Vortex silicon drift detector (Hitachi High-Technologies Science America, USA) equipped with a high – speed Xspress3 "readout" processing electronics (Quantum Detectors, UK), which was oriented at 90° to the incident beam and at 45° to the sample plane. Fluorescent energy windows were centered for As, Fe and other elements of interest (P, S, K, Ca, Zn, Cu, Mn). The samples were raster scanned in an oversampling regime (pixel size 35 μm x 35 μm) with beam dwell time 100 ms per pixel.

**Micro XFI and Micro-XAS of selected plant tissues**

Higher resolution imaging (2 –10 μm) of selected tissues of interest (areas of the apical root meristem of live main roots, and also selected areas of the 30 μm-thick cryo-sections of the apical shoot meristem) were performed at the bending magnet beamline 2-3 (SSRL) with the same geometric and fluorescent energy detection setup as described for the beamline 10-2 above. The beamline 2-3 is dedicated to micro-XFI and micro-XAS analysis of small samples. The beamline is equipped with a pair of Kirkpatrick-Baez (KB) mirrors as the focusing optics to achieve a spot size of 2 $\mu m^2$.

Pre-washed main roots were cut at approximately 0.5 cm from the root cap, so the cut part comprised the apical root meristem, the elongation zone and a part of maturation zone. The cut sections were placed on metal-free plastic coverslips (NuncThermanox, Thermo Scientific, USA), and covered with 4μm thick Ultralene ® XRF film. The samples of the shoot meristem with cotyledons were similarly prepared. The coverslips with specimens were fixed on the sample holders of the motorized stage and raster scanned with pixel size 10 μm × 10 μm (dwell time 25 ms) to get a coarse X-ray fluorescence image of the large area of the sample, and then the selected areas were scanned at the resolution 2 μm × 2 μm with dwell time 100 ms per pixel.

After high resolution images were obtained, we collected micro-XAS As-K near-edge spectra in the As 'hotspots' and other areas of interest. The XAS As-K near-edge spectra were analyzed using EXAFSPAK suite of programs, as described in the section XAS data analysis.

**Data Analysis for XFI (beamlines 10-2 and 2-3, SSRL)**

The beamlines 10-2 and 2-3 at the SSRL have a similar data collection setup. Data were collected with the MICROSCAN program with the XMPAGUI visual interface software. Fluorescence was normalized by the incident X-ray beam intensity I0 measured by the upstream ion chamber in each case. Data reduction was performed using custom Fortran-95 code, and included subtracting: (1) background signal from trace elements in the mount; and (2) the "tail" of the Compton scatter signal, determined from blank standard measurements. Concentrations of elements of interest (S, P, K, Ca, Cu, Zn, Mn, Fe, As) were converted to aerial concentrations ($\mu g/cm^2$) using reference standards (Micromatter, Vancouver, Canada) which were scanned with the same parameters as the samples. The elemental areal concentrations were calculated at the regions of interest (ROIs), using custom MATLAB (MathWorks ®) codes. The rendering of graphics was done with MicroAnalysis Toolkit software (Webb, 2011).

**X-ray Fluorescence Microscopy at the APS (2-ID-E station)**

X-ray fluorescence microscopy (XFM) was conducted at the 2-ID-E X-ray fluorescence microprobe station of the Advanced Photon Source (APS). APS is a 7 GeV synchrotron light source at Argonne National Laboratory. The 2-ID-E station is equipped with a Fresnel zone plate which allows focusing of the incident beam to a sub-micron spot size. The samples were prepared by sandwiching pieces of the live roots including the apical root meristem and elongation part between two 200 nm-thick low stress silicon nitride windows (Norcada ®, SiN window dimension 1.5 × 1.5 mm), fixed on motorized stage, as shown in Supplemental Figure 13, and raster-scanned with the incident beam energy 13.45 KeV. At first, "fly" scans with pixel size 2 μm × 2 μm and dwell time/pixel 30 ms were performed on a large area of the sample, after that a higher resolution 1 μm × 1 μm "fly" scans with dwell time per pixel 100 ms were conducted on the smaller area of the root tip, which included root cap and apical root meristem (Supplemental Figure 13).

The fluorescence radiation excited at each scan position was collected by an energy dispersive silicon drift detector (Vortex EM, SII Nanotechnology, Northridge, California, USA). Data were fitted and quantified by using two NBS thin film standards 1832 and 1833 (NIST, Gaithersburg, MD, USA). The fitting of two-dimensional elemental maps and analysis of the regions of interest (ROI) analysis were performed using MAPS software (Nietzold et al, 2018).

**Arsenic speciation in plant samples**

**Bulk XAS at the beamline 7-3 (SSRL)**

**Sample preparation**

The harvested plants were washed in deionized water, dissected on shoot and root parts, packaged in separate pre-cleaned Lucite cells, and secured using a layer of a self-adhesive metal-free Mylar tape. For calibration of arsenic concentration in the bulk XAS measurements, we used 10 μM and 50 μM solution of sodium meta-arsenite in deionized water in similar Lucite cells. The prepared samples and solutions were flash-frozen and stored in the liquid nitrogen environment prior the bulk XAS measurements. In a similar way, we produced samples of freshly prepared and solidified growth medium on agar base with 10 μM of sodium meta-arsenite added.

**Setup**

SSRL beamline 7-3 is dedicated to X-ray Absorption Spectroscopy of biological dilute solutions of metalloproteins. The beamline 7-3 is powered by a 20-pole, 2-Tesla wiggler, with no focusing optics used. Harmonic rejection is achieved by detuning one monochromator crystal so that the measured intensity is 50% of the maximum. A Si(220) double crystal monochromator was used in our experiments. The samples were placed at 45° to the incident beam and were maintained at ~10 K using a helium flow cryostat (Oxford Instruments, Abingdon, Oxfordshire, UK). Incident and transmitted intensities were measured using nitrogen-filled ion chambers and As K-edge fluorescence radiation was monitored using a 30-element germanium detector (Canberra Inc, Meriden, CT, USA). The beam size was adjusted to the sample size and had dimensions of 1.5 mm (vertical) x 8 mm (horizontal). For the bulk XAS of frozen plant samples, the energy range (energy sweep) was scanned in 10 eV steps in the pre-edge (11747 eV – 11847 eV) and the post edge (11897 eV - 12030 eV) regions, and in 0.25 eV steps in the near edge region (11847 eV – 11897 eV), with a dwell time of 1 s per energy point, with 3 energy sweeps collected for each sample.

**Energy calibration standards**

For the energy calibration of incident synchrotron radiation for both microprobe and bulk XAS measurements, we used transmission foils prepared from powdered grey As (alpha-phase As) on a nitrocellulose base. The main inflection (second derivative) point in the transmission spectrum was assigned a value of 11867 eV.

**Arsenic chemical standards**

For chemical speciation, we used a large library of As-K near-edge X-ray absorption spectra collected using a set of methylated, thiolated inorganic and organic arsenicals involved in arsenic metabolism in biological systems (Table S4).

**XAS data analysis**

No smoothing or filtering operations were performed on the data for XAS chemical speciation analysis. All data were processed using EXAFSPAK data analysis software (George and Pickering, 2000). The background was removed using the "BACKSUB" subroutine in the EXAFSPAK program, using an algorithm to normalize X-ray absorption data to tabulated mass absorption coefficients similar to that described earlier (Weng et al., 2005). Near-edge spectra were normalized to obtain an edge jump magnitude of unity.

For determining the minimal number of the principal components and the pre-selection of the standards, we used the PCA ("principal component analysis") and TARGET ("target transformation") subroutines together with evaluation of the factor indicator function from EXAFSPAK, based at the algorithms developed initially by Malinowski and Howery (1980). After pre-selection of the initial set of standards, we applied the DATFIT subroutine (least-square fitting using linear combination of chemical standards) to analyse chemical speciation in the samples (George and Pickering, 2000). A spectral component was removed if its contribution was less than either 1% of the total value or three times its estimated standard deviation. The results of the PCA analysis of the bulk As-K near-edge set shown in Supplemental Table 5, Table 6 and Supplemental Figure S12, indicate that just one component of the vector space would be sufficient to represent the spectra, with the best candidate represented by arsenic-tris-glutathione As(GS)3 standard (the least residual in the target transform test as listed in Supplemental Table 6). Similar results were obtained for the micro-XAS data (As-K near-edge spectra collected from hot spots in root apical meristem as shown in Figure 5E).

**Supplemental Table 1:** Identity matrix among plasma membrane residing PIN proteins (AtPIN1, AtPIN2, AtPIN3, AtPIN4, AtPIN7) from *Arabidopsis thaliana*

| | AtPIN1 | AtPIN2 | AtPIN3 | OsLsi2 |
|---|---|---|---|---|
| AtPIN1 | 100.00 | 62.09 | 66.00 | 15.38 |
| AtPIN2 | 62.09 | 100.00 | 60.95 | 15.96 |
| AtPIN3 | 66.00 | 60.95 | 100.00 | 16.77 |
| OsLsi2 | 15.38 | 15.96 | 16.77 | 100.00 |

**Supplemental Table 2.** Identity matrix of *Oryza sativa* silicon transporter (OsLsi2), *Arabidopsis thaliana* PINs (AtPIN1, AtPIN2, AtPIN3)

| | AtPIN1 | AtPIN2 | AtPIN3 | OsLsi2 |
|---|---|---|---|---|
| AtPIN1 | 100.00 | 62.09 | 66.00 | 15.38 |
| AtPIN2 | 62.09 | 100.00 | 60.95 | 15.96 |
| AtPIN3 | 66.00 | 60.95 | 100.00 | 16.77 |
| OsLsi2 | 15.38 | 15.96 | 16.77 | 100.00 |

**Supplemental Table 3.** Identity matrix of *Saccharomyces cerevisiae* ACR3 (SsACR3), *Pteris vittata* ACR3 (PvACR3) and *Escherichia coli* arsenite transporter (arsB) and *Oryza sativa* silicon transporter (OsLsi2)

| | ScACR3 | PvACR3 | arsB | OsLsi2 |
|---|---|---|---|---|
| ScACR3 | 100.00 | 40.87 | 10.89 | 9.27 |
| PvACR3 | 40.87 | 100.00 | 10.19 | 7.43 |
| arsB | 10.89 | 10.19 | 100.00 | 14.29 |
| OsLsi2 | 9.27 | 7.43 | 14.29 | 100.00 |

**Supplemental Table 4.** The library of As-K near-edge X-ray absorption fine structure spectra used for the data analysis.

| Abbreviation | Oxidation State | Semi-structural chemical formula | Chemical name(s) |
|---|---|---|---|
| As(GS)3 | As III | $[HO_2CCH(NH_2)CH_2CH_2CONHCH(CONHCH_2CO_2H)CH_2S\text{-}]_3$ As (oxidized) | arsenic-tris-glutathione; each GS group is glutathione (γ-glutamyl-cysteinyl-glycine) bound to As atom via its sulphur in cysteinyl residue |
| AsO4- | As V | [As(OH)2O2]- | arsenate in aqueous solution at ~pH7 |
| $As(OH)_3$ | As III | [As(OH)3] | arsenite in aqueous solution at pH~7 |
| MMAIII | As III | $[CH_3As(OH)_2]$ | monomethylarsonous acid at pH~7 |
| $MMA_{III}$-LIP | As III | $[CH_3As(S_2C_3H_5(CH_2)_4CO_2H]$ | monomethylarsonous residue bound to two sulphur atoms of the lipoic acid at pH~7; chemical name [ (2RS,4RS)-5-(2-methyl-1,3,2-dithiarsinan-4-yl) pentanoic acid ] |
| DMAIII | As III | $[(CH3)_2AsOH]$ | dimethylarsinous acid at pH ~7 |
| MMAV | As V | $[(CH3)As(OH)_2O\ ]$ | monomethylarsonic acid at pH~7 |
| DMAV | As V | [(CH3)2AsO2H] | dimethylarsenic acid pH~7 |
| TMAO | As V | $[(CH_3)_3As{=}O]$ | trimethyl arsine oxide |
| TMAS | As V | $[\ (CH3)_3As{=}S]$ | trimethylarsine sulfide |

**Supplemental Table 5.** Results of principal component analysis (PCA) of the set of bulk XAS As-K near-edge spectra (arsenite-exposed frozen roots and leaves of col-0 and *eir1-1*) represented by eigenvalues and IND function.

| Component | Eigenvalue | IND function × 10 $^{3}$ |
|---|---|---|
| 1 | 30.7540 | 0.893 |
| 2 | 0.1729 | 1.148 |
| 3 | 0.0262 | 6.221 |
| 4 | 0.0873 | 9999 |

**Supplemental Table 6**. Target transform results for reference standards in PCA analysis of bulk XAS As-K near-edge spectra (arsenite-exposed frozen roots and leaves of col-0 and *eir1-1*)

| Target reference standard | Residual × 10 $^{2}$ |
|---|---|
| As(GS)3 | 0.08679412 |
| MMAIII-LIP | 0.1095146 |
| DMAIII | 1.257795 |
| AsO4- | 4.409424 |
| MMAV | 4.464354 |
| MMAIII | 6.597711 |
| DMAV | 7.236827 |
| TMAO | 8.758961 |
| As(OH)3 | 12.66888 |
| TMAS | 13.5855 |

**Supplemental Figures**

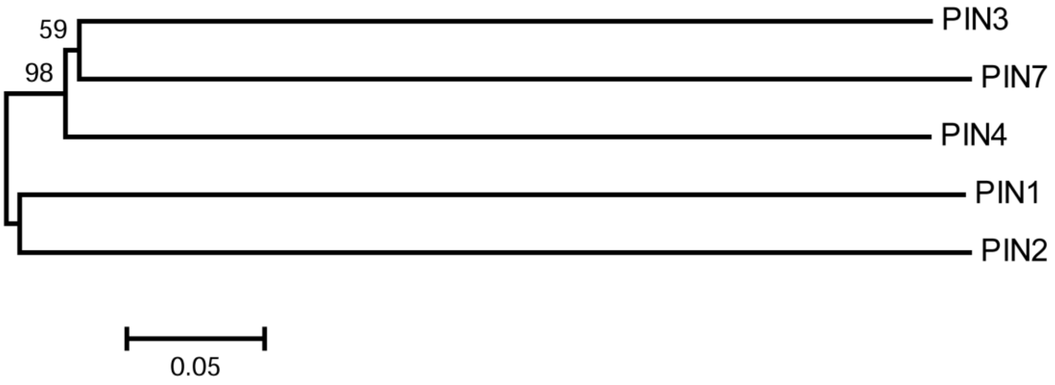


**Figure 1.** Cladogram of plasma membrane residing PIN proteins (AtPIN1, AtPIN2, AtPIN3, AtPIN4, AtPIN7) from *Arabidopsis thaliana* were constructed using MEGA6 (Molecular Evolutionary Genetics Analysis) software based on Neighbor Joining method and 1000 bootstrap test.

```
AtPIN1      MITAADFYHVMTAMVPLYVAMILAYGSVKWWKIFTPDQCSGINRFVALFAVPLLSFHFIA 60
AtPIN2      MITGKDMYDVLAAMVPLYVAMILAYGSVRWWGIFTPDQCSGINRFVAVFAVPLLSFHFIS 60
AtPIN3      MISWHDLYTVLTAVIPLYVAMILAYGSVRWWKIFSPDQCSGINRFVAIFAVPLLSFHFIS 60
AtPIN4      MITWHDLYTVLTAVVPLYVAMILAYGSVQWWKIFSPDQCSGINRFVAIFAVPLLSFHFIS 60
AtPIN7      MITWHDLYTVLTAVIPLYVAMILAYGSVRWWKIFSPDQCSGINRFVAIFAVPLLSFHFIS 60

AtPIN1      ANNPYAMNLRFLAADSLQKVIVLSLLFLWCKLSRNGSLDWTITLFSLSTLPNTLVMGIPL 120
AtPIN2      SNDPYAMNYHFLAADSLQKVVILAALFLWQAFSRRGSLEWMITLFSLSTLPNTLVMGIPL 120
AtPIN3      TNNPYAMNLRFIAADTLQKIIMLSLLVLWANFTRSGSLEWSITIFSLSTLPNTLVMGIPL 120
AtPIN4      TNDPYAMNFRFVAADTLQKIIMLVLLALWANLTKNGSLEWMITIFSLSTLPNTLVMGIPL 120
AtPIN7      SNNPYAMNLRFIAADTLQKLIMLTLLIIWANFTRSGSLEWSITIFSLSTLPNTLVMGIPL 120

AtPIN1      LKGMYGNFSGDLMVQIVVLQCIIWYTLMLFLFEYRGAKLLISEQFPDTAGSIVSIHVDSD 180
AtPIN2      LRAMYGDFSGNLMVQIVVLQSIIWYTLMLFLFEFRGAKLLISEQFPETAGSITSFRVDSD 180
AtPIN3      LIAMYGEYSGSLMVQIVVLQCIIWYTLLLFLFEFRGAKMLIMEQFPETAASIVSFKVESD 180
AtPIN4      LIAMYGTYAGSLMVQVVVLQCIIWYTLLLFLFEYRGAKLLIMEQFPETGASIVSFKVESD 180
AtPIN7      LIAMYGEYSGSLMVQIVVLQCIIWYTLLLFLFEYRGAKILIMEQFPETGASIVSFKVESD 180

AtPIN1      IMSLDGRQPLETEAEIKEDGKLHVTVRRSNASRSDIY------SRRSQGLSATPRPSNLT 234
AtPIN2      VISLNGREPLQTDAEIGDDGKLHVVVRRSSAASSMISSFNKSHGGGLNSSMITPRASNLT 240
AtPIN3      VVSLDGHDFLETDAEIGDDGKLHVTVRKSNASRRSFC------G-----PNMTPRPSNLT 229
AtPIN4      VVSLDGHDFLETDAEIGNDGKLHVTVRKSNASRRSL--------------MMTPRPSNLT 226
AtPIN7      VVSLDGHDFLETDAQIGDDGKLHVTVRKSNASRRSFY------GG--GGTNMTPRPSNLT 232

AtPIN1      NAEIYSLQSSRNPTPRGSSFNHTDFYSMMASGGGRNSN------------FGPGEA--VF 280
AtPIN2      GVEIYSVQSSREPTPRASSFNQTDFYAMFNASKAPSPRHGYTNSYGGAGAGPGGDVYSLQ 300
AtPIN3      GAEIYSLS----TTPRGSNFNHSDFYNMMGFPGGRLSN------------FGPADMYSVQ 273
AtPIN4      GAEIYSLS----STPRGSNFNHSDFYSVMGFPGGRLSN------------FGPADLYSVQ 270
AtPIN7      GAEIYSLN----TTPRGSNFNHSDFYSMMGFPGGRLSN------------FGPADMYSVQ 276

AtPIN1      GSKGPTPRPSNYEEDGGPAKPTAAGTAAGAGRFHYQSG-GSGGGGGAHYPAPNPGMFSPN 339
AtPIN2      SSKGVTPRTSNFDEEVMKTAKKAG-RGGRS-----MSGELYNNNSVPSYPPPNPMFTGST 354
AtPIN3      SSRGPTPRPSNFEENCAMASSP---------RFGYYPG-----GGAGSYPAPNPEFSSTT 319
AtPIN4      SSRGPTPRPSNFEENNAV-------------KYGFYNNTNSSVPAAGSYPAPNPEFSTGT 317
AtPIN7      SSRGPTPRPSNFEESCAMASSP---------RFGYYPG-----GAPGSYPAPNPEFSTGN 322

AtPIN1      TGGGGGTAAKG---------NAPVVGGKRQDGNGRDLHMFVWSSSASPVSDVFGGG---- 386
AtPIN2      SGASGVK-KK-------ESGGGGSGGGVGVGGQNKEMNMFVWSSSASPVSEANAKNAMTR 406
AtPIN3      TSTANKSVNKNPKDVNTNQQTTLPTGGKSNSHDAKELHMFVWSSNGSPVSDRAGLNVFGG 379
AtPIN4      GVS------TKPNKIPKENQQQLQEKDSKASHDAKELHMFVWSSSASPVSDVFG----GG 367
AtPIN7      KTG-----SKAPKE-------NHHHVGKSNSNDAKELHMFVWGSNGSPVSDRAGLQVDNG 370

AtPIN1      GGNHHADYSTATNDHQKDVKISVPQG----------NSNDNQYVEREEFSFGNKDDDSKV 436
AtPIN2      GSSTD---------VSTDPKVSIPPHDNLA-TKAMQNLIENMSPG-------RKGHVE-- 447
AtPIN3      APDNDQ--GGRSDQGAKEIRMLVPDQSHNGETKAVAHPASGDFGGEQQFSFAGKEEEAER 437
AtPIN4      AGDNVA--TEQSEQGAKEIRMVVSDQPRKSNARGGGDDI----GGL------DSGEGERE 415
AtPIN7      ANE-QV--GKSDQGGAKEIRMLISDHTQNGENKA--GPMNGDYGGE---------EESER 416

AtPIN1      LAT--D---------------GGNNISNKTTQAKVMPPTSVMTRLILIMVWRKLIRNPNS 479
AtPIN2      --MDQDGNNGGK-SPYMGKKGSDVEDGGPGPRKQQMPPASVMTRLILIMVWRKLIRNPNT 504
AtPIN3      PKDAENGLNKLAPNSTAALQSKTGLGGAEASQRKNMPPASVMTRLILIMVWRKLIRNPNT 497
AtPIN4      IEKATAGLNKMGSNSTAELEAAGGDGGG--NNGTHMPPTSVMTRLILIMVWRKLIRNPNT 473
AtPIN7      VKEVPNGLHKLRCNSTAELNPKEAIETGETVPVKHMPPASVMTRLILIMVWRKLIRNPNT 476

AtPIN1      YSSLFGITWSLISFKWNIEMPALIAKSISILSDAGLGMAMFSLGLFMALNPRIIACGNRR 539
AtPIN2      YSSLFGLAWSLVSFKWNIKMPTIMSGSISILSDAGLGMAMFSLGLFMALQPKIIACGKSV 564
AtPIN3      YSSLIGLIWALVAFRWHVAMPKIIQQSISILSDAGLGMAMFSLGLFMALQPKLIACGNSV 557
AtPIN4      YSSLIGLIWALVAYRWHVAMPKILQQSISILSDAGLGMAMFSLGLFMALQPKIIACGNSV 533
AtPIN7      YSSLIGLIWALVAFRWDVAMPKIIQQSISILSDAGLGMAMFSLGLFMALQPKLIACGNST 536

AtPIN1      AAFAAAMRFVVGPAVMLVASYAVGLRGVLLHVAIIQAALPQGIVPFVFAKEYNVHPDILS 599
AtPIN2      AGFAMAVRFLTGPAVIAATSIAIGIRGDLLHIAIVQAALPQGIVPFVFAKEYNVHPDILS 624
AtPIN3      ATFAMAVRFLTGPAVMAVAAIAIGLRGDLLRVAIVQAALPQGIVPFVFAKEYNVHPAILS 617
AtPIN4      ATFAMAVRFITGPAIMAVAGIAIGLHGDLLRIAIVQAALPQGIVPFVFAKEYNVHPTILS 593
AtPIN7      ATFAMAVRFFTGPAVMAVAAMAIGLRGDLLRVAIVQAALPQGIVPFVFAKEYNVHPAILS 596

AtPIN1      TAVIFGMLIALPITLLYYILLGL 622
AtPIN2      TAVIFGMLVALPVTVLYYVLLGL 647
AtPIN3      TGVIFGMLIALPITLVYYILLGL 640
AtPIN4      TGVIFGMLIALPITLVYYILLGL 616
AtPIN7      TGVIFGMLIALPITLVYYILLGL 619
```

**Figure 2.** Multiple Sequence Alignment of plasma membrane residing PIN proteins (AtPIN1, AtPIN2, AtPIN3, AtPIN4, AtPIN7) from *Arabidopsis thaliana*

```
AtPIN2     ----------------------------------------MITGKDMYDVLAAMVPLYVA 20
SsACR3     -----------MSEDQK-SENSVPSKVNMVNRTDILTTIKSLSWLDLMLPFTIILSIIIA 48
PvACR3     MENSSAERKQQLALDIADGNDPSDAAKNPDGRTKLQGLFKQLSLLDRYLYVWIFIVMAVS 60
arsB       ----------------------------------------------MLLAGAIFVLTIVL 14
AtPIN1     ----------------------------------------MITAADFYHVMTAMVPLYVA 20
AtPIN3     ----------------------------------------MISWHDLYTVLTAVIPLYVA 20

AtPIN2     MILAY---GSVRWWGIFTPDQCSGINRFVAVFAVPLLSFHFI------SSNDPYAMNYHF 71
SsACR3     VIISVYVPSSRHTFDAEGHPNLMGV-------SIPLTVGM-IVMMIPPICKVSWESIHKY 100
PvACR3     IIFGYYVKGVKKAFQV---AEITSV-------SLPIAIGL-WVMMYPVLCKVQYEILGGV 109
arsB       VIWQ--PKGLGIGWSA-----TLGA-------VLALISGVVHIGDIPVVWNIVWNATATF 60
AtPIN1     MILAY---GSVKWWKIFTPDQCSGINRFVALFAVPLLSFHFI------AANNPYAMNLRF 71
AtPIN3     MILAY---GSVRWWKIFSPDQCSGINRFVAIFAVPLLSFHFI------STNNPYAMNLRF 71

AtPIN2     LAADSLQKVVILAALFLWQAFSRR------GS----------LEWMITLFSLSTL-PNTL 110
SsACR3     FYRSYIRKQLALSLFLNWVIGP----------LLMT-----ALAWM-ALFDYKEYRQGII 144
PvACR3     LRQAGSLKTISLSVVLNWVVGP----------ALMT-----GLAWA-TLPDLPDFRTGVI 153
arsB       IAVIIISLLLDESGFFEWAALHVSRWGNGRGRLLFTYIVLLGAAVA-ALFAN----DGAA 115
AtPIN1     LAADSLQKVIVLSLLFLWCKLSRN------GS----------LDWTITLFSLSTL-PNTL 114
AtPIN3     IAADTLQKIIMLSLLVLWANFTRS------GS----------LEWSITIFSLSTL-PNTL 114

AtPIN2     VMGIPLLRAMY---GDFSG-------------NL----MVQIVVLQSIIWYTLMLFLFEF 154
SsACR3     MIGVARCIAMVLIWNQIAGG----------DNDLCVVLVITNSLLQMVLYAPLQIFY--- 191
PvACR3     LVGIARCIAMVLIWNDLAKG----------DADYCAILVAINSILQIILFTPVALLYL-- 201
arsB       LILTPIVIAMLLALGFSKGTTLAFVMAAGFIADTASLPLIVSNLV--------------- 160
AtPIN1     VMGIPLLKGMY---GNFSG-------------DL----MVQIVVLQCIIWYTLMLFLF-- 152
AtPIN3     VMGIPLLIAMY---GEYSG-------------SL----MVQIVVLQCIIWYTLLLFLFEF 154

AtPIN2     -----------------ISI-----LSDAGLGMAMFSLGLFMALQPK----IIACGKS-- 563
SsACR3     IFISRG-YQFIHEIGSAILCFVPLMLYSCIAWFLTFALMRYLSISRSDTQRECSCDQELL 323
PvACR3     MFSIQA-HQIVDNIGHVVRVAVPLLLYFGILFFGSLGICRWLK----------------- 312
arsB       VVYGLRNAGLTEYLSGVLNV-----LADKGLWAATFGTGFLTAFLSS----IMNNMPT-- 340
AtPIN1     --------EMPALIAKSISI-----LSDAGLGMAMFSLGLFMALNPR----IIACGNR-- 538
AtPIN3     --------AMPKIIQQSISI-----LSDAGLGMAMFSLGLFMALQPK----LIACGNS-- 556

AtPIN2     ----------VAG--------FAMAVRFLTGPAVIAATSIA------IGIR-GDLLHIAI 598
SsACR3     LKRVWGRKSCEASFSITMTQCFTMASNNFELSLAIAISLYGNNSKQAIAATFGPLLEVPI 383
PvACR3     -----------VPYPLMVTQCFTAASNNFELAIAVAVGSFGTDSTQALAATIGPLIEVPV 361
arsB       ----------V-----------------LVGALSI-DGSTA------TGVIKEAMIFANV 366
AtPIN1     ----------RAA--------FAAAMRFVVGPAVMLVASYA------VGLR-GVLLHVAI 573
AtPIN3     ----------VAT--------FAMAVRFLTGPAVMAVAAIA------IGLR-GDLLRVAI 591

AtPIN2     VQAALPQGIVPFVFAKEYNVHPDILSTAVIFGM-----L--VALPVTVLYYVL------- 644
SsACR3     LL-----------------------ILA-------------IVARILKPYYIWTNRN--- 404
PvACR3     LL-----------------------LFV-------------YIVGFFQRK----GPSV-- 379
arsB       IG---------------CDLGPKITPIGSLATLLWLHVLSQKNMTITWGYYFRTGIVMTL 411
AtPIN1     IQAALPQGIVPFVFAKEYNVHPDILSTAVIFGM-----L--IALPITLLYYILLGL---- 622
AtPIN3     VQAALPQGIVPFVFAKEYNVHPAILSTGVIFGM-----L--IALPITLVYYILLGL---- 640
```

**Figure 3.** Homology among PIN proteins of *Arabidopsis thaliana and Saccharomyces cerevisiae* ACR3 (SsACR3), *Pteris vittata* ACR3 (PvACR3) and *Escherichia coli* arsenite transporter (ArsB).

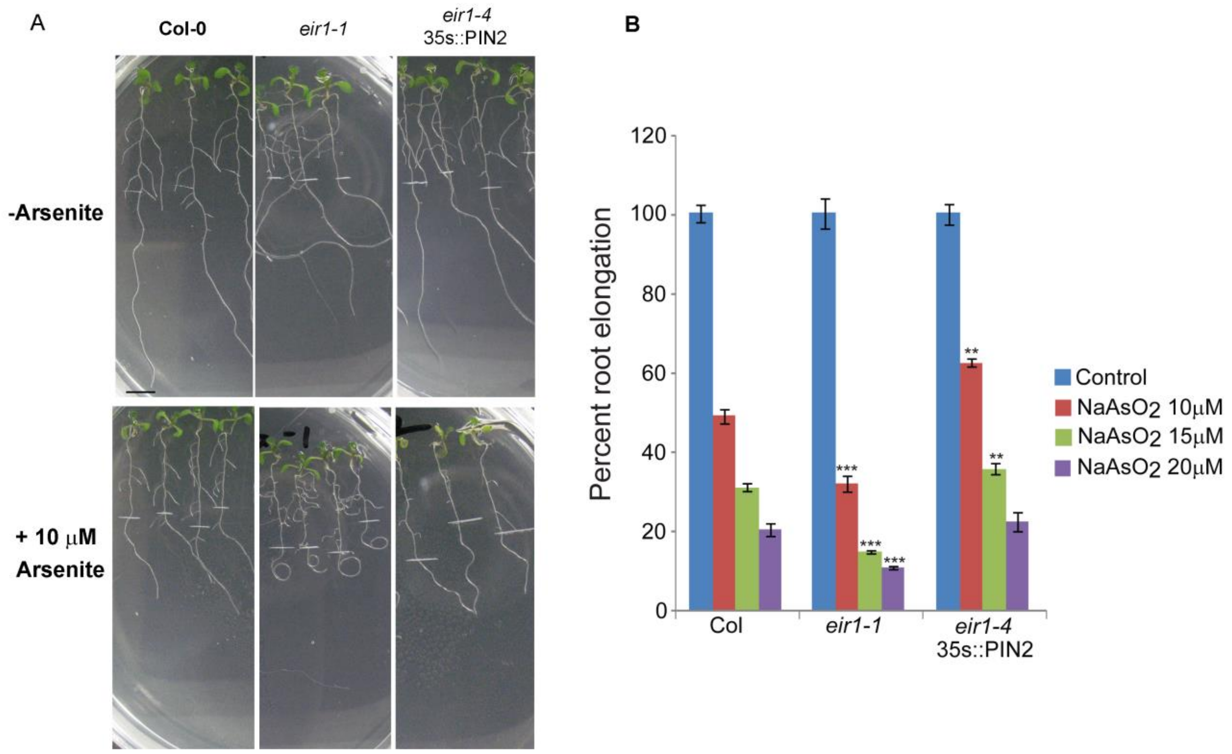


**Figure 4. Effect of arsenite on root elongation response of *eir1-1* and PIN2 overexpression line**

Five-day-old light grown wild-type or mutants seedlings were transferred to new agar plates supplemented with or without arsenite and incubated for three days under continuous light. **(A)** Representative images of root phenotype of wild-type, *eir1-1* and *PIN2* overexpression line after 10 μM arsenite treatment for 3 days. Bar represents 0.5cm. Please note the curly root phenotype of *eir1-1* in response to arsenite treatment. **(B)** Root elongation response in presence of various concentration of arsenite after 3-day incubation. Overexpression of PIN2 results in resistance to arsenite-induced root growth inhibition as judged by Student's *t*-test ($P<0.001$).

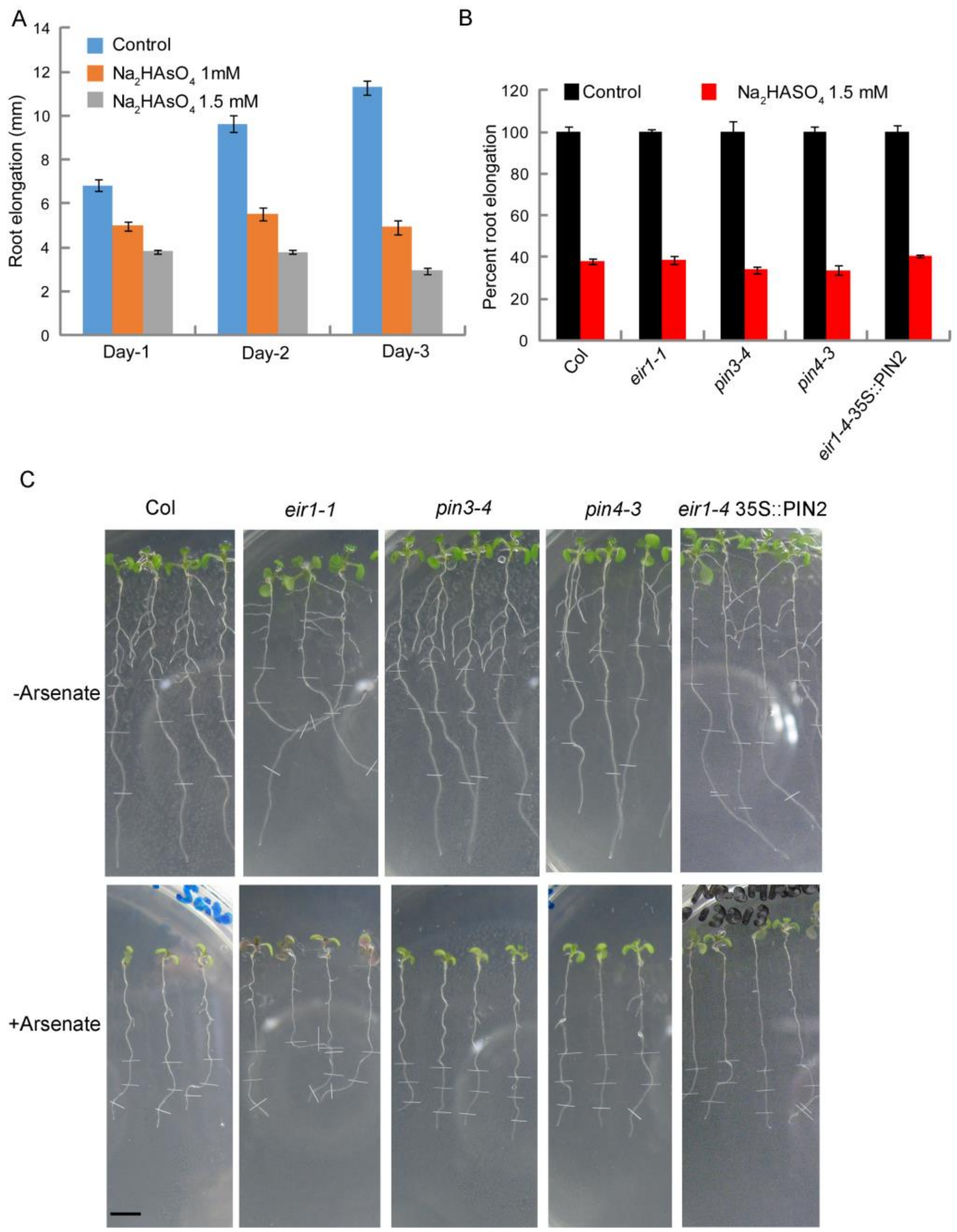


**Figure 5. Effect of arsenate on wild-type and mutants root elongation response**

Five-day-old light grown wild-type or mutants seedlings were transferred to new agar plates supplemented with or without arsenate and incubated for various time lengths under continuous light. **(A)** Time course of arsenate induced inhibition of root elongation in wild-type. **(B)** Root elongation response of pin mutants in presence of arsenate after 3-day incubation. Approximately fifty percent inhibition of root growth was observed at 10μM arsenate. **(C)** Representative images of root phenotype of wild-type, *pin* mutants and *pin2* complemented line after 1.5mM arsenate treatment for 3 days. Bar represents 0.5cm.

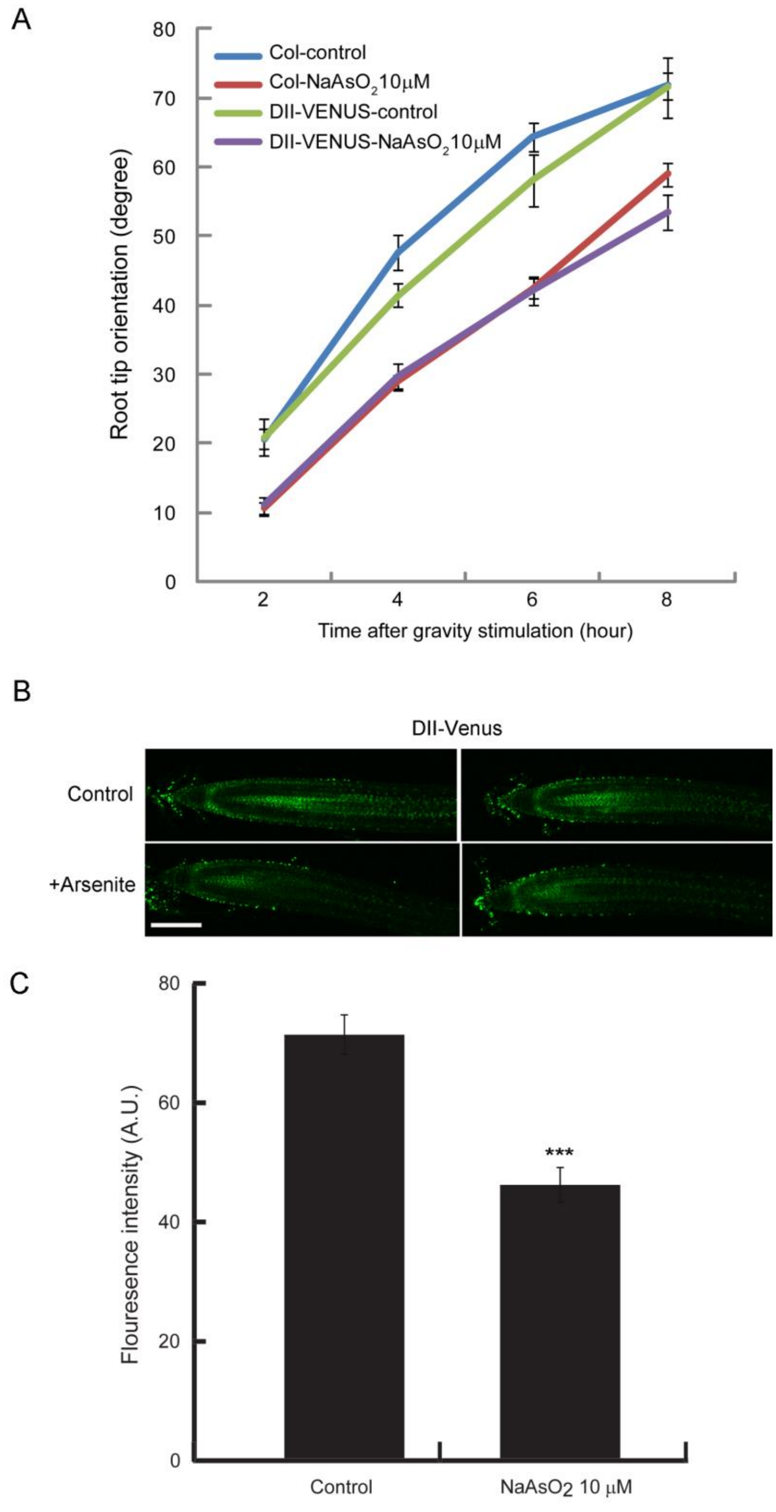


**Figure 6. Arsenite inhibits root gravity and alters intracellular auxin response.**

(A) Effect of Arsenite on root gravity response.

For assaying gravitropism, five –day-old light grown seedlings were transferred to arsenite, and gravity stimulation was provided to the root by rotating the plates 90°. Data for root tip orientation was collected for 2,4,6 and 8 h. Vertical bars represent mean ± S.E. of the experimental means from at least five independent experiments ($n$ = 5 or more), where experimental means were obtained from 8-10 seedlings per experiment. Arsenite-induced inhibition of root gravity response was significant for both genotypes at all time point as judged by Student's $t$-test ($P < 0.0001$).

**(B)** Long term effect of Arsenite on intracellular auxin response

Five-day-old light grown DII-VENUS seedlings were transferred to new agar plates and subjected to arsenite incubation for 3 days and imaged with confocal microscopy using the same settings. Note the reduced DII-VENUS signal in arsenite-treated roots at all-time points. Bar represents 100 □m. **(C)** Quantification of fluorescence intensities. Vertical bars represent mean ± S.E. of the experimental means from three independent experiments ($n$ = 3), where experimental means were obtained from 8-10 seedlings per experiment. Compared with the control treatment, arsenite-induced reduction of DII-VENUS signal was highly significant ($P < 0.0001$) as judged by Student's $t$-test. A.U., arbitrary unit.

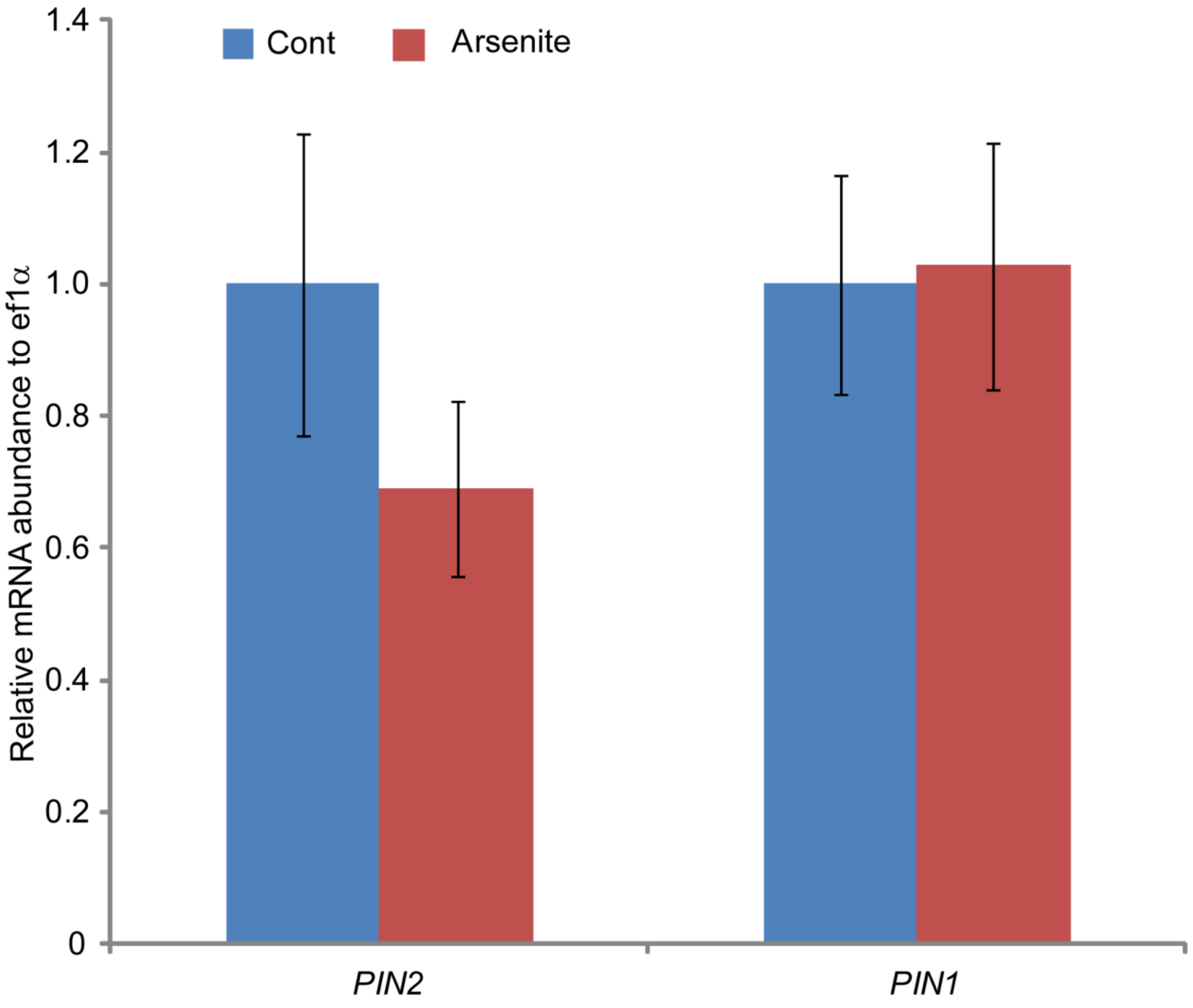


**Figure 7. Arsenite does not alter the expression of *PIN* genes.**
Five-day-old light grown seedlings were transferred to new agar plates and subjected arsenite treatment for 3 days under continuous light condition. *PIN*s expression was analyzed in the root of Arabidopsis seedlings using qRT-PCR. The data were obtained from 3 independent biological replicates with three technical replicates for each biological replicate. Effect of arsenite on PIN2 and PIN1 gene expression was statistically insignificant as judged by Student's *t*-test ($P > 0.30$ for PIN2 and $P > 0.90$ for PIN1).

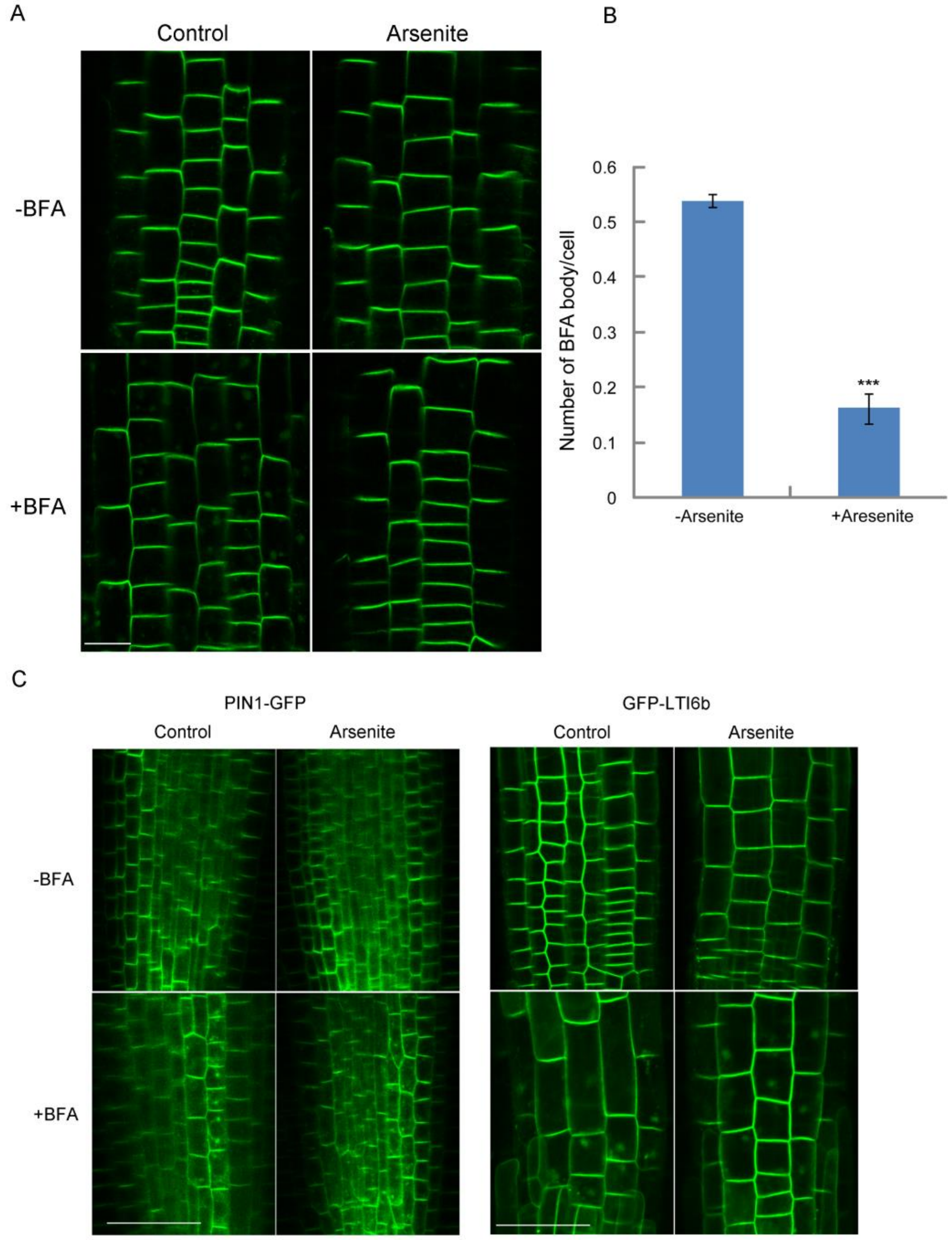


**Figure 8. Effect of long term treatment of Arsenite on intracellular protein trafficking** 5-day-old PIN2::PIN2–GFP, PIN1::PIN1-GFP and EGFP-LTI6b transgenic seedlings were treated with arsenite for 3d. After the incubation, seedlings were treated with 20 μM BFA for 30 min. The images were captured using same confocal setting and are representative of 15-20 roots obtained from at least 4 independent experiments.

**(A)** Effect of arsenite on PIN2 trafficking. Bar represents 10 μm. **(B)** Quantitative analysis of formation of PIN2-BFA body in the transition zone of PIN2::PIN2–GFP transgenic plants in presence or absence of arsenite. Both arsenite treated and control seedlings were treated with 20 μM BFA for 40 min and subjected to imaging. The images were captured using same confocal setting and area and total number of BFA body and number of cells were counted in the imaged area. Bar graph represents the average number of BFA body formed per cell. Vertical bars represent mean ± S.E. of the experimental means from at least four independent experiments ($n$ = 4 or more), where experimental means were obtained from 5-6 seedling images per experiment. **(C)** Effect of arsenite on PIN1 and LTI6b trafficking. Note that BFA bodies are formed in presence of arsenite. Bar represents 50 μm.

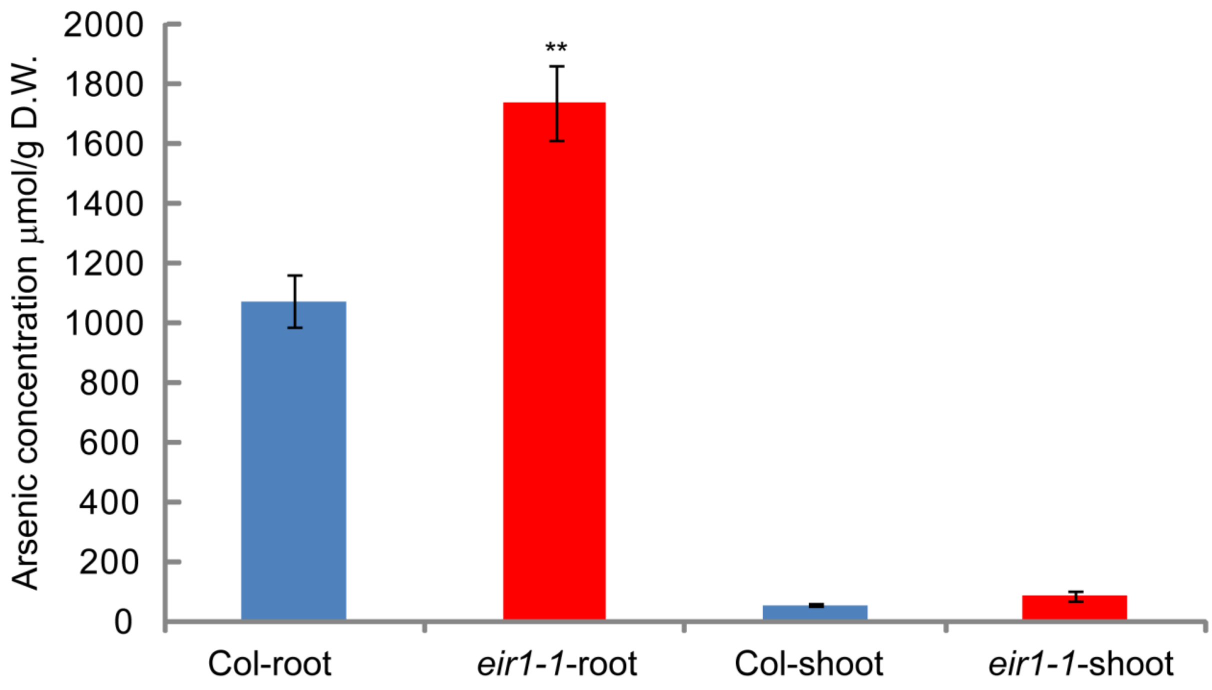


**Figure 9. Increased accumulation of arsenic in *eir1-1/pin2* mutant after long term exogenous treatment with arsenite.**

Five-day-old seedlings were subjected to 10µM arsenite treatment for 3 day. After the incubation arsenic content was measured as described in methods. The experiments were conducted using at least three biological replicates. For each biological replicate, three technical replicates were assayed (20-25 seedlings were used per sample). Asterisks represent the statistical significance between genotypes (** $P < 0.001$).

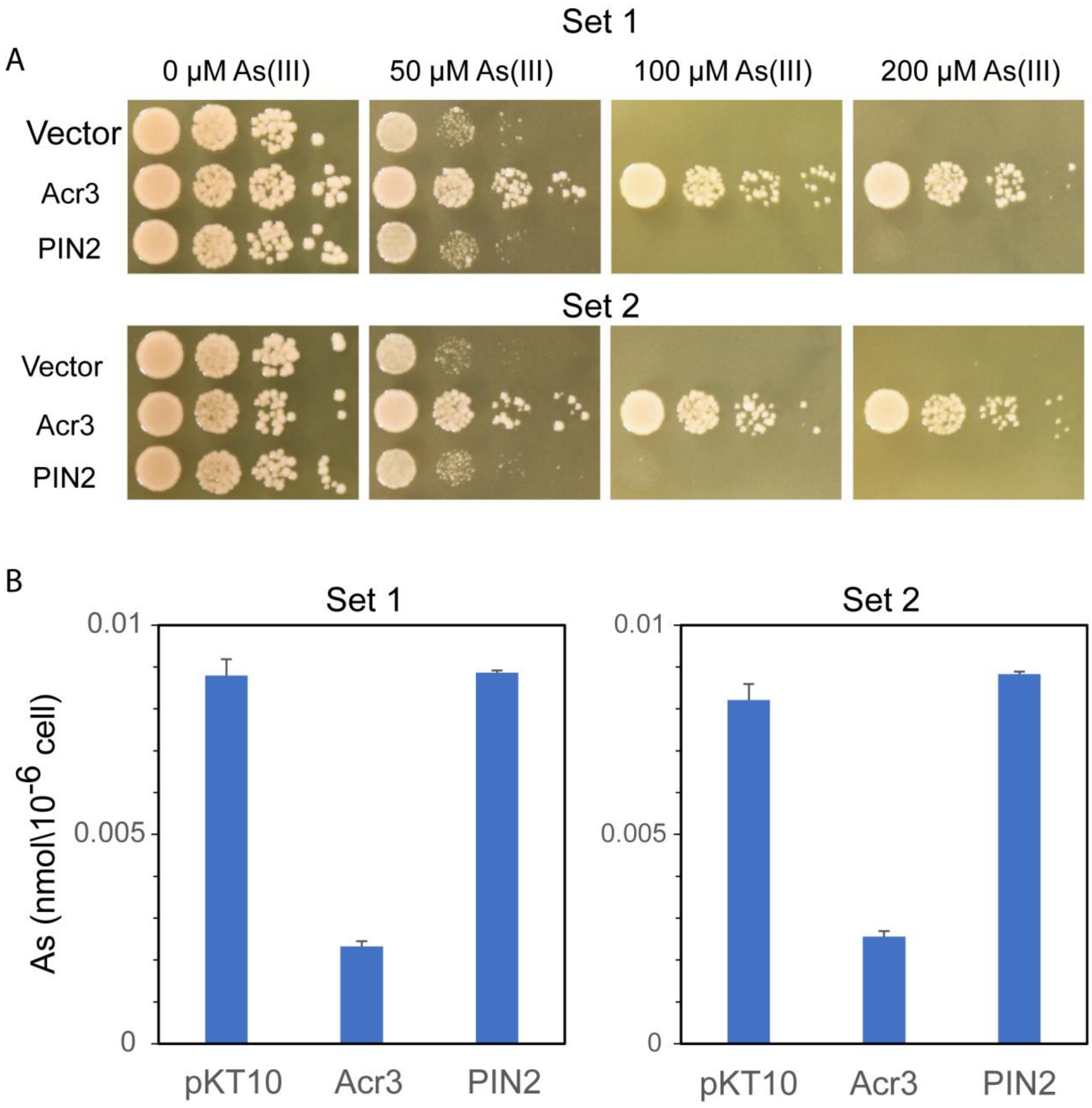


**Figure 10. PIN2 is incapable of transporting Arsenite in *Saccharomyces cerevisiae***
(A) As (III) tolerance assay. Two independent clones (Set 1 and 2) were spotted onto YPD medium containing As(III) and photos were taken after 48 hrs. Each spots were sequentially diluted by 10 times from left to right. **(B)** As (III) uptake experiments. The yeast carrying the plasmids were incubated with SD medium containing 100 μM As (III) for 1 h and applied for As (III) determination by ICP-MS. The data represent mean ± SD (n=4).

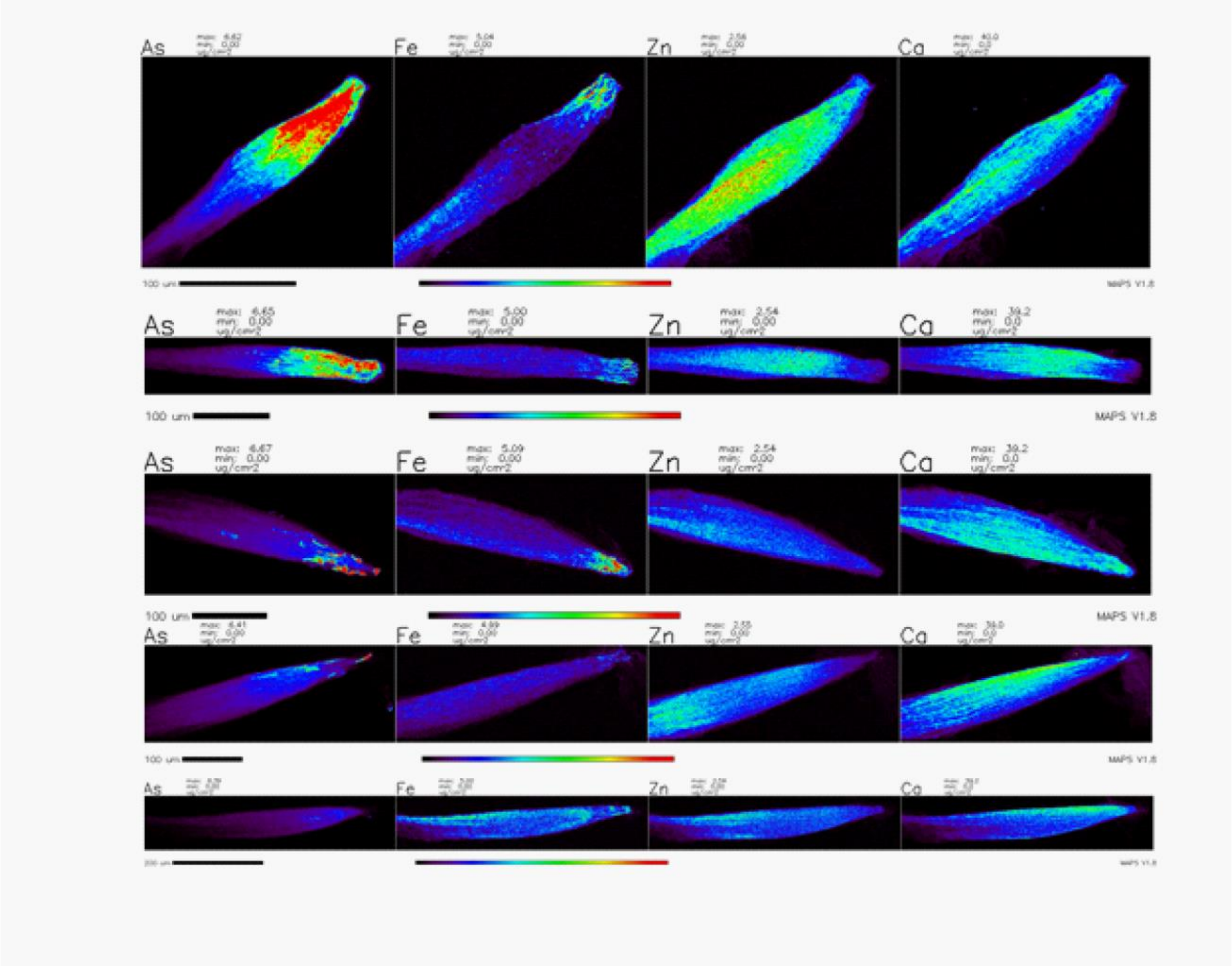


**Figure 11**. **Areal densities (µg/cm$^2$) of As, Fe, Zn, Ca in the apical meristem of roots of Arabidopsis.** The plants were exposed to 10 µM sodium meta-arsenite for 3 d in the growth medium. The top two rows- *pin2* mutant , the bottom three rows -Col-0. Brighter colors denote higher values of areal densities. (APS, 2-ID-E, scanned with step 2 µm, dwell time per pixel 100 ms).

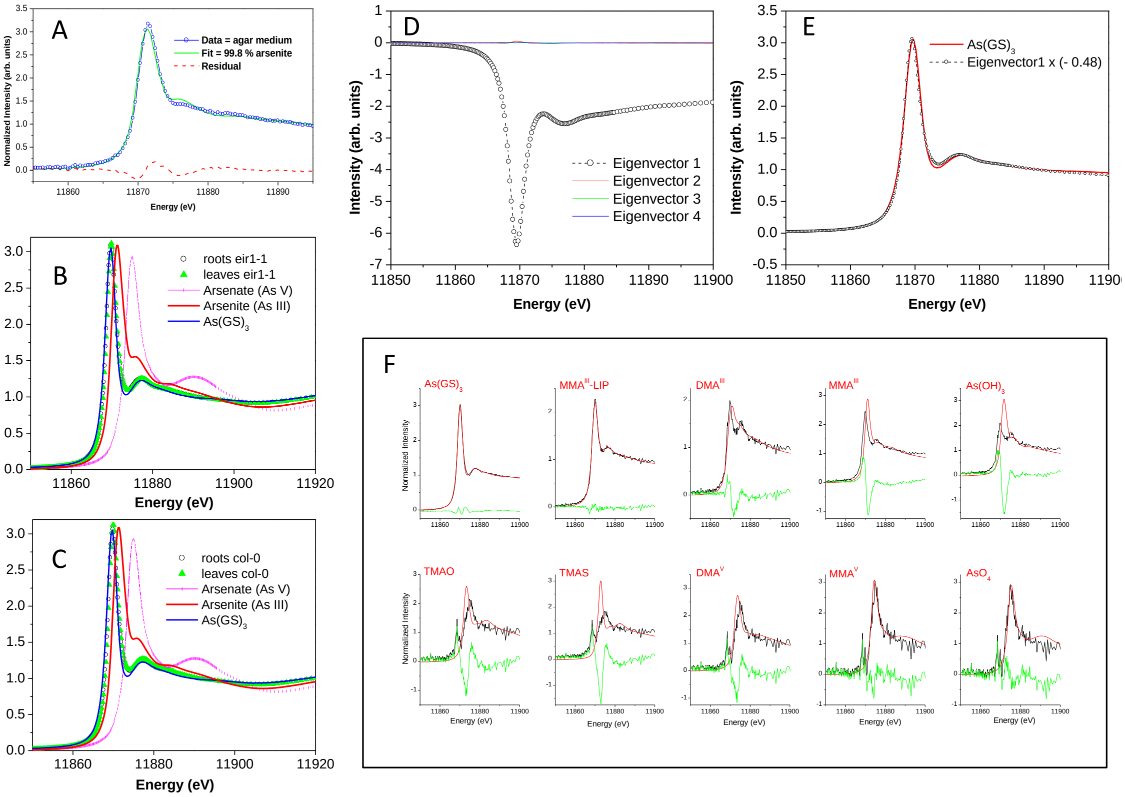


**Figure 12. Arsenic speciation analysis (bulk As-K near-edge XAS). (A)** Least square fitting to the linear combination of standard spectra for the near edge As-K XAS of fresh agar-based growth medium with 10 µM arsenite shows that mixing and solidification of agar does not affect arsenic form in the original medium. This spectrum is best fitted with arsenite in aqueous solution at pH ~ 7 ($AsO_4^-$) standard. The data are marked by a blue line with empty circles, the fitting curve – by a green solid line and the residual of the fit is denoted by a dashed red line. **(B)** Bulk near-edge As-K XAS of flash frozen leaves (denoted by green triangles) and roots (circles) of eir1-1 plants harvested after exposure to 10 µM sodium meta-arsenite from $5^{th}$ to $8^{th}$ DAG in the growth medium are almost identical to XAS spectrum of arsenic-tris-glutathione $As(GS)_3$ standard. The spectra of arsenite (As III) and arsenate (As V) in aqueous solution pH ~ 7 are shown for comparison. **(C)** The same as **(B)**, for leaves and roots of col-0 plants. **(D)** The first 4 eigenvectors from PCA analysis of bulk As-K XAS spectra collected from frozen leaves and roots of arsenite-exposed col-0 and eir1-1 plants. **(E)** The first eigenvector from **(D)** scaled by factor (-0.48) and plotted side-a-side with arsenic-tris-glutathione $As(GS)_3$ standard. **(F)** Target transform test for the library of arsenical standards listed in **Table S2**, for the vector space obtained using PCA analysis of the bulk near-edge As-K XAS of leaves and roots. The best fit with a smallest residual corresponds to arsenic-tris-glutathione $As(GS)_3$ standard.

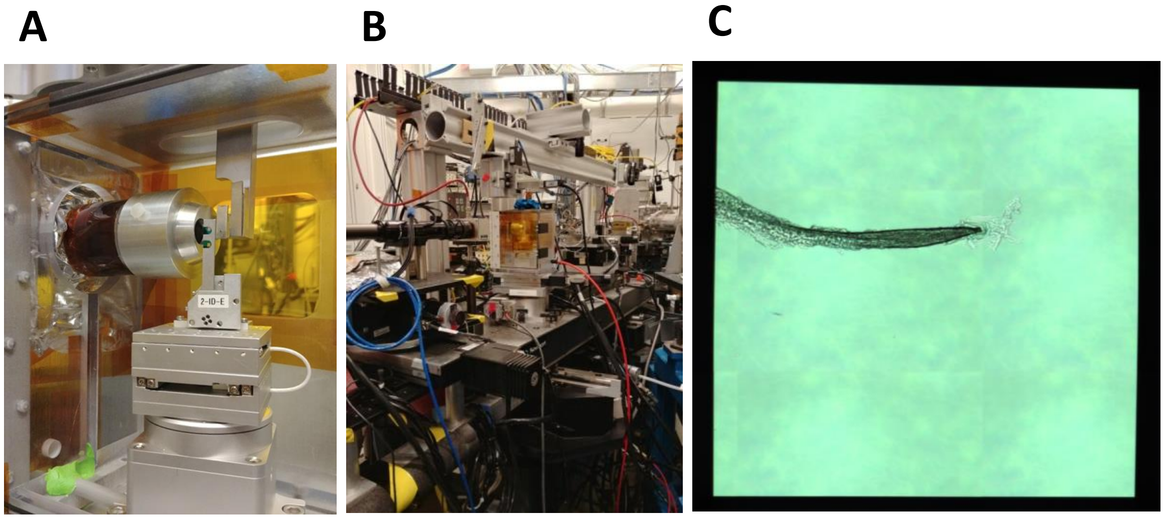


**Figure 13. XFI setup at the beamline 2-ID-E at the APS**.
**(A)** Silicon nitride windows with mounted hydrated root samples are glued to the aluminum stick holder and placed on the motorized stage with detector axis oriented at 45° to the sample plane. **(B)** A view of the beamline 2-ID-E hatch with the sample stage chamber in the center. **(C)** A digital image of a hydrated untreated col-0 root tip mounted between 2 silicon nitride windows.
